# SOCS1 expression in prostate epithelial cells is essential for tissue homeostasis and tumor suppression

**DOI:** 10.64898/2026.05.09.723770

**Authors:** Awais Ullah Ihsan, Mozhdeh Namvarpour, Mohammad Moradzad, Anny Armas Cayarga, Evelyne Ng Kwan Lim, Diya Binoy Joseph, Stephanie Petkiwicz, Eric Massé, Akihiko Yoshimura, Gerardo Ferbeyre, Alfredo Menendez, Sheela Ramanathan, Subburaj Ilangumaran

**Affiliations:** Departments of Immunology and cell biology, Biochemistry and functional genomics Université de Sherbrooke, Sherbrooke J1H 5N4, Quebec, Canada, Institut de recherche sur le cancer de l’Université de Sherbrooke (IRCUS), Sherbrooke, QC J1E 4K8, Canada; Departments of Immunology and cell biology, Faculty of Medicine and Health Sciences, Université de Sherbrooke, Sherbrooke J1H 5N4, Quebec, Canada; Institute for Stem Cell Science and Regenerative Medicine (inStem), Bengaluru 560065, Karnataka, India; Department of Pathology and Laboratory Medicine, The Ottawa Hospital and the University of Ottawa, Ottawa, Canada; Department of Microbiology and Immunology, Keio University School of Medicine, Tokyo 160-8582, Japan; Department of Biochemistry and Molecular medicine, Montréal H3C 3J7, Quebec, Canada; Department of Microbiology and infectious diseases, Faculty of Medicine and Health Sciences, Université de Sherbrooke, Sherbrooke, QC, J1H 5N4, Canada, Institut de recherche sur le cancer de l’Université de Sherbrooke (IRCUS), Sherbrooke, QC J1E 4K8, Canada

**Author notes:** **Corresponding authors:** Subburaj Ilangumaran, PhD, Sheela Ramanathan, PhD, Faculty of Medicine and Health Sciences Université de Sherbrooke, 3001, 12^th^ avenue North Sherbrooke, QC J1H 5N4, Canada. Equal contribution (Awais Ullah Ihsan, Mozhdeh Namvarpour).

**Keywords:** Prostate gland, SOCS1, hyperplasia, high-fat diet, UPEC, colibactin, prostate cancer, proteomics

## Abstract

Suppressor of cytokine signaling 1 (SOCS1) negative regulates inflammatory cytokine production and attenuates oncogenic growth factor signaling pathways. Reduced SOCS1 protein expression in human prostate cancer correlates with greater disease severity. To define the physiological functions of SOCS1 functions in the prostate, we conditionally ablated *Socs1* in prostate epithelial cells of C57BL/6 mice. These *Socs1^ΔPE^* mice exhibited normal prostate development, maturation and lobular architecture. However, adult *Socs1^ΔPE^*mice developed progressive epithelial hyperplasia and inflammatory cell infiltration that were temporally and spatially distinct. SOCS1-deficient prostate showed increased epithelial cell proliferation and elevated oxidative stress markers, and prostate organoids recapitulated this hyperplasia phenotype. Diet-induced obesity exacerbated both hyperplasia and inflammation in SOCS1-deficient prostate. Upon transurethral infection with uropathogenic *Escherichia coli* UPEC1677 expressing the genotoxin colibactin, *Socs1^ΔPE^* mice developed invasive prostate cancer with complete loss of lobular architecture, whereas control mice developed hyperplasia and pre-neoplastic lesions. *In vitro*, SOCS1-deficient prostate organoid–derived epithelial cells exhibited increased DNA damage following exposure to UPEC1677. Deletion of the colibactin biosynthetic gene *clbP* in UPEC1677 abolished its ability to induce DNA damage in SOCS1-deficient cells and to drive prostate cancer *in vivo*. Proteomic analysis of prostate organoids revealed dysregulation of basal and luminal epithelial lineage markers and signaling pathway proteins that could promote neoplasia in SOCS1-deficient cells. Collectively, these findings establish an essential, epithelial cell-intrinsic role for SOCS1 in maintaining prostate tissue homeostasis by restraining proliferation, regulating lineage plasticity, limiting inflammation and oxidative stress, and conferring protection against genotoxic injury and neoplastic transformation.

## Introduction

Prostate cancer is a leading cause of cancer mortality among ageing men ^1^. Inflammatory lesions are frequently observed in prostate biopsy specimens, and epidemiological evidence suggests that chronic inflammation increases the risk of both benign prostatic hyperplasia (BPH) and prostate cancer ^2,3^. Prostate cancer often coexists with BPH, and the two conditions share certain genetic alterations ^4^. Although BPH is considered a potential risk factor for prostate cancer, a definitive causal relationship has not been established ^5,6^.

Ethnicity and family history are established risk factors for prostate cancer. However, fewer than 5% of the genetic alterations identified in prostate cancer are attributable to inherited susceptibility ^7^. Cancer development is multifactorial, involving both intrinsic, non-modifiable genetic alterations and extrinsic, potentially modifiable risk factors. Given that prostate cancer predominantly arises in older men, cumulative exposure to environmental factors, particularly those that promote prostatic inflammation, likely contribute to prostate cancer pathogenesis ^7–9^. Such non-intrinsic risk factors include unhealthy diet and obesity, urogenital infections and potential carcinogens ^3,10,11^. Many of these factors are thought to induce chronic inflammation, leading to the production of reactive oxygen and nitrogen species that can cause DNA damage and drive genetic and epigenetic alterations in caretaker genes, tumor suppressors, and oncogenes ^7,8,12^. Consistent with this notion, several inflammation-related and DNA mismatch repair genes have been associated with prostate cancer ^7,13^. Oncogenic alterations such as *NKX3.1* downregulation, *MYC* amplification, *TEMPRSS2-ERG* gene fusion and *PTEN* inactivation are associated with distinct stages of prostate cancer development and progression ^14^. Even though the functional roles of these lesions have been demonstrated in genetically engineered mouse models, the timing of their occurrence during tumorigenesis and their causal relationship to disease initiation remain to be fully resolved ^14,15^.

Inflammation of the prostate tissue is believed to initiate a cycle of tissue injury, proliferative inflammatory atrophy (PIA) and regenerative responses that can progress to BPH, prostatic intraepithelial neoplasia (PIN), and ultimately prostate cancer ^16^. Chronic prostatitis is poorly understood in molecular terms despite its clinical relevance to prostate cancer ^2^. In inflammatory settings, pro-inflammatory cytokines, chemokines and growth factors produced by stromal and immune cells support tissue repair. However, their excessive production and deregulated signaling can perpetuate inflammation and foster cancer initiation and progression. For example, IL-6 and TNFα are implicated in promoting both prostatitis and proliferation of prostate cancer cell lines ^17,18^, and transgenic expression of IL-6 in the mouse prostate gland induces spontaneous cancer development ^19^.

Several regulatory mechanisms control inflammatory cytokine signaling to maintain tissue homeostasis ^20^. Among these regulatory factors, suppressor of cytokine signaling 1 (SOCS1) was identified as a negative feedback regulator of IL-6-induced JAK-STAT signaling in macrophages. However, genetic ablation of the *Socs1* gene in mouse revealed that SOCS1 is largely dispensable for attenuating IL-6 signaling but is essential for restraining IFNγ signaling ^21^. In addition, SOCS1 controls inflammatory responses by attenuating lipopolysaccharide (LPS)-induced TLR4 signaling in macrophages, thereby limiting the production of inflammatory cytokines such as IL-6 and TNFα ^22,23^. Beyond cytokine pathways, SOCS1 also inhibits growth factor induced receptor tyrosine kinase signaling pathways ^24,25^.

SOCS1 is widely regarded as a tumor suppressor as its expression is frequently silenced by CpG methylation in diverse cancers, and its overexpression inhibits tumor growth in xenograft models ^25^. In human prostate cancer cell lines, the *SOCS1* gene is induced by IL-6 and androgens, and a peptide corresponding to its kinase inhibitory region suppresses IL-6-driven proliferation ^26,27^. Moreover, elevated expression of *miR-30d*, which reduces SOCS1 transcript levels, is significantly associated with prostate cancer recurrence ^28^, while upregulation of SOCS2, which promotes SOCS1 degradation, correlates with increased malignancy ^29^. SOCS1 also attenuates HGF-induced MET signaling in prostate cancer cell lines, limiting proliferation and migration ^30^. In clinical specimens, SOCS1 protein expression decreases with increasing tumor grade and metastasis ^31^. Collectively, these findings strongly suggest a tumor suppressor role for SOCS1 in the prostate. To directly test this hypothesis and to identify SOCS1-regulated oncogenic signaling pathways in prostate cancer, we genetically ablated the *Socs1* gene in prostate epithelial cells of C57Bl/6 mice. Our findings on these *Socs1^ΔPE^* mice show that SOCS1 loss disrupts tissues homeostasis in the prostate gland leading to age-dependent hyperplasia and synergize with genotoxic insults to cause prostate cancer.

## Materials and Methods

### Mice

*Socs1^fl/fl^* mice were backcrossed to C57BL/6 mice obtained from Charles River Laboratories for more than ten generations ^32^. *Pb-Cre4* (B6.D2-Tg(Pbsn-cre)4Prb) mice ^33^ were purchased from Mouse Models of Human Cancer Consortium (MMHCC) of the National Cancer Institute. As the *Pb-Cre4* transgene is expressed in oocytes, only male *Pb-Cre4* mice were crossed with *Socs1^fl/fl^* females to generate *Socs1^fl/fl^Pb-Cre4* male offspring, lacking SOCS1 specifically in prostate epithelial cells (*Socs1^ΔPE^*). Mice were housed in ventilated cages with 12 hours day-night cycle and fed with normal chow *ad libitum*. All experiments on mice were carried out during daytime with the approval of the Université de Sherbrooke Ethics Committee for Animal Care and Use (Protocol ID: 2020-2502 and 2024-4410).

### Histology

The internal urogenital system was removed in entirety and placed in a Petri dish. After removing the seminal vesicles, urethra and the urinary bladder, the prostate was pulled out, surrounding adipose tissues gently teased out and positioned in a histology cassette to reveal all four lobes of the prostate upon sectioning, following consensus guidelines ^34^ (**Supplementary Figure S1**). The cassette was fixed in 4% paraformaldehyde (PFA) overnight and stored in 70% ethanol before embedding in paraffin blocks. Formalin-fixed paraffin-embedded (FFPE) sections of prostate tissues (5 µm) were deparaffinized in xylene, gradually rehydrated in graded alcohol, and stained with hematoxylin and eosin (H&E). Digital images of stained sections were acquired using a Nanozoomer Digital Pathology (NDP) slide scanner (Hamamatsu Photonics, Japan) and the images were analyzed using the NDP.view 2.7.52 software (Hamamatsu Photonics).

### Immunofluorescence

Prostate sections were deparaffinized, rehydrated, and antigenic epitopes were retrieved using the Antigen Retriever (Electron microscopy Sciences, Hatfield, PA). After cooling, the sections were incubated in blocking buffer (5% BSA in PBS containing 0.1% Triton-X) for 60 min at room temperature. Primary antibodies (**Supplementary Table S1**) diluted in blocking buffer were applied and incubated overnight at 4°C. After washing, slides were incubated with secondary antibodies conjugated to Alexafluor-488 or Alexafluor-568 fluorochromes (**Supplementary Table S1**) at room temperature for 60 min. The slides were washed, incubated with DNA staining dye DAPI (ThermoFisher; Cat# 62248) in PBS for 5 min, washed and mounted with coverslips using DAKO Fluorescence mounting medium (DAKO, Cat# S3023). After allowing the slides to dry for 2 h, images were acquired using Axioskop 2 fluorescence microscope (Carl Zeiss Canada Ltd, Toronto, Canada).

### Immunohistochemistry

Endogenous peroxidase activity in tissue sections was blocked after antigen retrieval by incubating in 3% hydrogen peroxide for 10 min. After washing and incubation with blocking buffer and primary antibody, slides were washed and incubated with SignalStain Boost IHC detection reagent conjugated to horseradish peroxidase antibody (HRP, Mouse; Cell Signaling Technology, Cat# 8125) for 1 hour at room temperature. After washing, slides were treated with 3,3’-diaminobenzidine substrate (Vector Laboratories, Cat# SK-4100) for 5-8 min at room temperature. Sections were counterstained with hematoxylin for 10 sec, washed, dehydrated, and mounted with Fischer Chemical^TM^ Permount^TM^ mounting medium (Fischer Scientific; Cat# SP15-100). Slides were allowed to dry 24 h before scanning using the Nanozoomer.

### Prostate organoid cultures

Prostate epithelial organoid cultures were established following published methods ^35^. Briefly, prostate glands from 5-6-month-old mice were removed and minced into small pieces, digested in serum-free Gibco^TM^ Advanced DMEM/F12 medium (ThermoFischer Scientific, Cat# 12634-010) containing 4 mg/mL of Collagenase II (Worthington, Cat# LS004176) and 10 μM of Y-27632/ROCK inhibitor (Stem Cell Technologies, Cat# 72304) in a 15 mL tube for 1.5 hours at 37°C with gentle intermittent pipetting. The digested tissue was centrifuged at 400 *g* for 5 min at 4°C. The pellet was suspended in 2 mL of Gibco^TM^ TrypLE (ThermoFischer Scientific, Cat# 12605-010) and incubated at 37°C for 30 min with intermittent pipetting. The tissue digest was centrifuged at 400 *g* for 5 min, and cells were resuspended in AdDMEM/F12 and passed through a Falcon^®^ 40 μm cell strainer (Corning, Cat# 352340) to clear the debris. The solution was centrifuged at 400 *g* for 5 min and cell pellets were resuspended and counted. Cells were sedimented and resuspended in Matrigel^®^ (Corning, Cat# 356255), liquified by placing on ice, and 20,000 cells were placed as a 40 μL drop in each well of a 24-well plate. The plate was placed in the 37°C incubator for 15 min to allow the Matrigel to polymerize. Pre-warmed AdDMEM/F12 containing glutaMAX 20 μL/mL (Gibco^TM^ ThermoFisher Scientific, Cat# 35050-061), HEPES, 20 mM (Millipore Sigma, Cat# H0887), penicillin-streptomycin, 100 units/mL (Wisent, Cat# 450-201-EL), B-27^TM^, 20 μL/mL (Gibco^TM^ ThermoFisher Scientific, Cat# 12587-010), N-acetyl cysteine (1.25 mM, Millipore Sigma, Cat# A-9165), mouse epidermal growth factor (50 ng/mL; PeproTech Cat# 315-09), mouse Noggin, 50 μL/mL (STEMCELL Technologies, Cat# 78061.2), R-Spondin, 50 μL/mL (R&D Systems^TM^ ThermoFisher Scientific, Cat# 7150RS050), TGFβ pathway inhibitor A83-01 (50 nM, STEMCELL Technologies, Cat# 72022), 5α-dihydrotestosterone (1 nM, Sigma Aldrich, Cat# A8380) and the ROCK inhibitor Y-27632 dihydrochloride (10 μM, STEMCELL Technologies, Cat# 72304) was added to the wells and the culture plate returned to the incubator. Organoid growth was monitored every day and media was changed on alternate days. Brightfield images of organoids were taken on days 3, 5 and 8 to monitor organoid growth.

### Processing organoids for histology

Fully grown organoids were collected on day 8 and processed for histology. Briefly, media was aspirated and chilled AdDMEM/F12 was added and the Matrigel was gently disrupted with a pipette tip to aid its dissolution. The contents were transferred to Eppendorf tubes and centrifuged at 300 *g* for 4 min at 4°C. The pellets were suspended in 4% PFA for 30 minutes, washed and resuspended in 70% ethanol and stored at 4°C overnight. The organoids were pelleted down, and resuspended in 30 µL of pre-warmed Eperdia^TM^ Histogel^TM^ (ThermoFisher Scientific, Cat# HG-4000-012) to encapsulate the organoids, transferred to a mold on a cold block for 30 min. The solidified Histogel containing organoids was extruded into a histology cassette, transferred to 70% ethanol and processed for paraffinization, blocking and sectioning. The organoid sections transferred to slides were processed for histochemical and immunofluorescence staining following the same methods used for prostate tissue sections.

### Organoid formation efficiency

After establishing organoid cultures, the number of live organoids in each Matrigel were counted seven days later when stable morphology was attained. Because of the three-dimensional nature of the organoid culture, counting was performed directly under bright-field microscope (4× magnification) at different depths of field. Organoid formation efficiency was calculated as the number of organoids counted over the number of cells seeded (organoid formation count, OFC). Additionally, organoids were classified based on morphological features as ‘luminal’ (distinct lumen) or compact (with no or smaller lumen). Independent biological replicates were derived from different animals and at least three replicates per sample were counted.

### Genotoxic bacteria

A clinical isolate of uropathogenic *Escherichia coli,* UPEC1677 ^36^, was obtained from Dr. Alexander de Vos (Center for Experimental & Molecular Medicine, Amsterdam University Medical Centers) and grown in Luria-Bertani (LB) broth. The *clbP* gene in UPEC1677 was deleted by homologous recombination to disrupt the colibactin biosynthetic pathway. Briefly, a PCR reaction on pKD4 plasmid (Addgene, Cat# 45605) was performed using primers listed in **Supplementary Table 2** to introduce sequences homologous to *clbP* flanking the kanamycin resistance cassette. The PCR product was electroporated into UPEC1677 carrying the pSIM6 plasmid ^37^, followed by the induction of the λ Red system at 42°C for 15 minutes to delete *clbP*. Kanamycin-resistant colonies of UPEC1677*ΔclbP* were selected and *clbP* deletion was verified by sequencing using primers listed in **Supplementary Table S2** (**Supplementary Figure S2**). Log phase cultures of UPEC and UPEC1677 *ΔclbP* (OD600 ∼0.5) were centrifuged and bacteria resuspended at 2× 10^8^ colony forming units (CFU) per mL.

### Transurethral bacterial inoculation

Male *Socs1^fl/fl^* and *Socs1^ΔPE^* mice were infected with UPEC1677 or UPEC1677*ΔclbP* at 12-16 weeks of age by transurethral route under isofluorane anesthesia following published methods ^38^. Briefly, the penile body of anesthetized mouse was exposed and a sterile 2.5 inch-long polyethylene tubing (BD Intramedic PE10; Fisher Scientific, Cat# 22204008; outer diameter 0.61 mm), lubricated with Vaseline and attached to the smoothened tip of a 27G needle (Terumo, Cat# KL0325) and containing 10 µL of bacterial inoculum (2×10^6^ CFU), was inserted into the urethral opening and guided to the prostate. After bacterial instillation, mice were kept on heating pads under anesthesia for 30 min to prevent urine voiding and bacterial loss. Mice were housed in sterile cages and monitored for discomfort and mortality. Most mice tolerated the procedure except a few that died within 1-2 weeks. The mice were routinely monitored over several months. When more than one UPEC1677-instilled mouse developed discomfort and malaise, which occurred 6-7 months post-inoculation, all mice were euthanized and their prostate glands examined. Their prostate tissues were embedded in paraffin for histological examination. DNA damage was assessed using immunofluorescence staining for phosphorylated histone γH2AX ^39^.

### Infection of prostate epithelial organoid cells with UPEC1677

Prostate epithelial organoids from SOCS1-deficient and control mice were dissociated into single cells using mechanical and enzymatic disruption. Briefly, organoids were washed with ice-cold DMEM, mechanically disrupted by pipetting with cut 1 mL pipette tips, and pelleted by centrifugation. The pellet was resuspended in TrypLE and incubated at 37 °C for 5–10 min with gentle pipetting until a single-cell suspension was obtained. Enzymatic activity was stopped with complete medium, and cells were counted and seeded on sterile glass coverslips in 12-well plates at ∼10^5^ cells per well in DMEM supplemented with 10% FBS and standard additives. Cells were cultured for 48 h to form confluent epithelial monolayers. In parallel, UPEC1677 and UPEC1677*ΔclbP* were cultured on LB agar to obtain single colonies and expanded overnight in LB broth. On the day of infection, bacteria were grown to mid-log phase (OD600 ∼1.0) and washed in PBS containing calcium and magnesium. The bacterial inoculum was prepared in serum-free DMEM and added to organoid-derived epithelial monolayers at the multiplicity of infection (MOI) of approximately 80 based on estimated cell numbers at the time of infection (∼2.5 × 10⁵ cells per well). After incubation at 37°C in a CO2 incubator, the monolayers were washed thoroughly and incubated in DMEM containing 10 %FBS and gentamicin at 50 µg per mL for 1.5 hours to eliminate extracellular bacteria, then maintained in medium containing 5 µg/mL gentamicin for 16 to 20 h. Cells were subsequently washed, fixed with 4% PFA overnight at 4°C. DNA damage was assessed by immunofluorescence detection of γH2AX as a marker of double strand DNA breaks.

### Quantification of fluorescently stained cells

Ki67 and γH2AX-positive cells were quantified by using NIH ImageJ (version 1.8.0_172, https://imagej.nih.gov/ij/download.html). Immunofluorescence images from randomly selected fields were analyzed by splitting channels, and nuclei were identified based on DAPI staining. A consistent threshold was applied across all images within each experiment to define positive staining. The total number of cells were then calculated using “analyze particles” option, and fluorescent cells were counted manually and divided by total number of cells to achieve percentage of fluorescence positive cells per field and were averaged for each group.

### Statistical Analysis

Statistical analyses and graph generation were performed using GraphPad Prism (version 10.6; San Diego, CA, USA).

### Organoid mass spectrometry

Prostate epithelial organoids established from *Socs1^fl/fl^*and *Socs1^ΔPE^* mice were collected on day 8. Cells were isolated, proteins were extracted and digested with trypsin, and peptides were collected as detailed in Supplementary methods. Liquid chromatography-tandem mass spectrometry (LC-MS/MS) analysis of peptides, protein identification and proteomic data analysis are described in Supplementary methods. Significantly modulated proteins were identified using the cut-off values of fold change (FC) ≤-1 and ≥1 and adjusted *p*-value (False discovery rate, FDR) <0.05 and <0.1.

## Results

### SOCS1 is essential to maintain homeostasis in the post-pubertal prostate gland

*Socs1^ΔPE^* mice are viable, healthy and fertile, and their prostate glands did not reveal signs of any gross morphological abnormality compared to those of *Socs1^fl/fl^* littermate controls for up to thirteen months of age. However, histological examination of the prostate from 9-12-month-old mice revealed notable changes in the *Socs1^ΔPE^* mouse prostate (**Fig. 1A-B**). Whereas the glandular acini of control *Socs1^fl/fl^* mouse prostates displayed a single layer of epithelium in all four lobes, with papillary patterns in the anterior lobe and moderate infoldings in anterior and dorsal lobes, *Socs1^ΔPE^* mouse prostates frequently displayed extensive tufting in anterior and dorsal lobes with focal areas of multilayered epithelium, and patches of epithelial cell accumulation in lateral lobes, indicating epithelial hyperplasia (**Fig. 1A-B**). Even though ventral lobes of *Socs1^ΔPE^* mice harbored single layered epithelia, they displayed variable degrees of peri-glandular inflammatory cell infiltration (**Fig. 1B**). The hyperplastic prostate epithelia of *Socs1^ΔPE^* mice did not display nuclear atypia and the basement membranes of glandular acini were intact. Because the prostate continues to grow after puberty under the influence of androgens ^40,41^, we examined younger *Socs1^ΔPE^* mice (**Supplementary Fig. S3A-B**). Even though the probasin gene promoter is active and the Pb-Cre transgene is expressed in the prostate one week after birth ^33^, *Socs1^ΔPE^* mice did not show any histologically detectable changes in the prostate up to three months of age, indicating that SOCS1 deficiency does not affect prostate development or maturation. However, nearly a third of *Socs1^ΔPE^* mice developed discernible changes in epithelial cell layers of anterior, dorsal and lateral prostate lobes at 3-5 months of age. Cumulative data show that the incidence of prostate hyperplasia increased upon ageing, reaching 60% at 8 months and attained 100% penetrance by 12 months (**Fig. 1C**). In comparison, *Socs1^fl/fl^* mice displayed did not show any hyperplastic changes until 9 months of age, with only 25% of mice developing hyperplasia of much lower magnitude at 12 months of age (**Fig. 1C**). Inflammatory infiltrates in the ventral prostate lobes of *Socs1^ΔPE^*mice did not appear until 6 months, however, all *Socs1^ΔPE^* mice displayed inflammatory cell infiltration by 12 months of age (**Fig. 1D**). Very few *Socs1^fl/fl^* mice showed inflammatory cell infiltration of the prostate at 8 months, and their proportion did not increase further. Overall, the progressive increase in incidence and severity of prostate hyperplasia and inflammatory cell infiltration in *Socs1^ΔPE^*mice compared to *Socs1^fl/fl^* controls indicate that SOCS1 plays an essential non-redundant role in maintaining homeostasis of the prostate gland while attenuating inflammatory responses. Moreover, the temporal and spatial dissociation of hyperplasia and inflammatory cell infiltration in different prostate lobes of *Socs1^ΔPE^* mice suggest diverse regulatory functions of SOCS1 in prostate epithelial cells in anatomically segregated prostate lobes.

**Figure 1.**
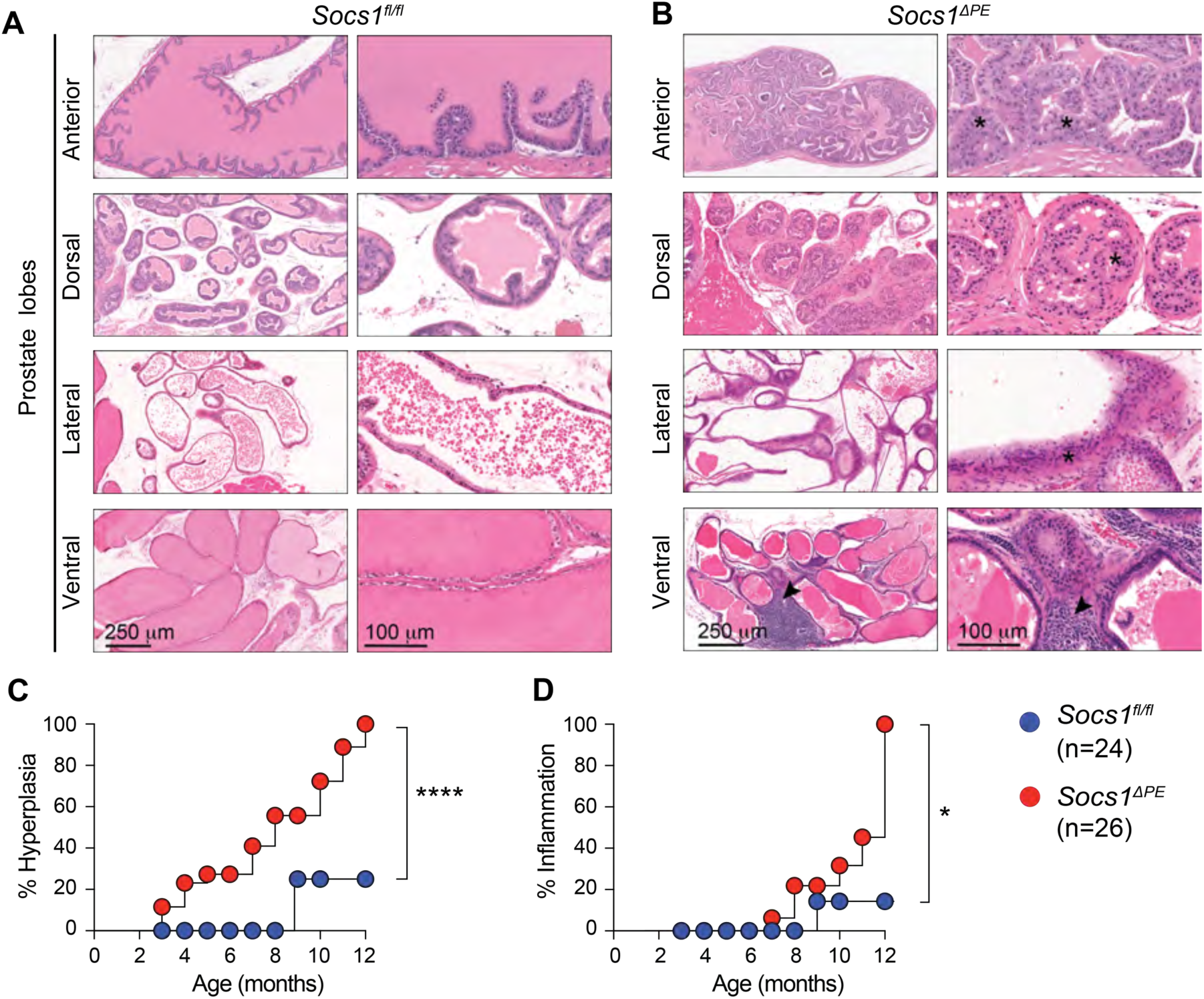
SOCS1 is critical to maintain homeostasis in post-pubertal prostate gland. (A,B) FFPE-sections of prostate glands from 9-12 months old *Socs1^fl/fl^* (A) and *Socs1^ΔPE^* (B) mice were stained with hematoxylin and eosin, and images of entire prostate glands captured using Nanozoomer digital pathology scanner as shown in Supplementary Figure S1. Representative regions of anterior, dorsal, lateral and ventral prostate lobes are shown. Hyperplastic prostate epithelium in anterior and dorsal lobes of *Socs1^ΔPE^* mice is indicated by asterisks, and inflammatory cel infiltration in ventral lobes by arrowheads. Sections of prostate glands from 3-5 and 6-8 months-old mice are shown in Supplementary Figure S3. (C,D) Cumulative data on the incidence of hyperplastic features, characterized by multilayered luminal epithelium, prominently displayed in anterior and dorsal lobes (C), and inflammatory cell infiltration, often occurring in ventral lobes (D). Data were collected from 6, 8 and 10 *Socs1^fl/fl^*mice, and 8, 9 and 9 *Socs1^ΔPE^* mice from the age groups of 3-5, 6-8 and 9-12 months. Data were statistically compared using the Log-rank (Mantel-Cox) test. *p* values: * <0.05; **** <0.0001.

### SOCS1 loss promotes epithelial cell proliferation in the adult prostate gland

Signaling pathways induced by cytokines and growth factors such as IL-6 and HGF, which are implicated in prostate epithelial cell proliferation, can be regulated by SOCS1 ^26,27,30,42–45^. Therefore, we evaluated cell proliferation in the prostate lobes of 6-8 months-old *Socs1^ΔPE^* and control mice by Ki67 staining. Anterior and dorsal prostate lobes of *Socs1^ΔPE^* mice contained significantly more Ki67+ cells than those of *Socs1^fl/fl^* mice (**Fig. 2A-B**). This trend also occurred in the prostate glands of 3-5 months-old mice and continued until 12 months age (**Fig. 2B, Supplementary Fig. S4**). Lateral lobes of *Socs1^ΔPE^* mice did not show increase in cell proliferation, whereas ventral lobes contained more Ki67+ staining in the epithelial layer in the 9-12 months age group, although some Ki67+ staining also occurred within the peri-glandular regions containing inflammatory cells (**Fig. 2B**, **Supplementary Fig. S4**). The distribution of basal and luminal epithelial cells, detected using CK5 and CK8 markers, respectively, showed a normal distribution in all prostate lobes of *Socs1^ΔPE^* mice as in control *Socs1^fl/fl^*mice (**Fig. 2C**). This pattern of epithelial cell distribution remained intact until 12 months of age in *Socs1^ΔPE^* mice (**Supplementary Figure S5**). However, occasional CK8+CK5+ cells expressing both luminal and basal cell markers, which are considered intermediate or bipotent progenitor cells ^46,47^, were identifiable in anterior and dorsal prostate lobes of 3-5 and 6-8 months-old *Socs1^ΔPE^* mice but not in *Socs1^fl/fl^* controls (**Fig. 2C**, **Supplementary Fig. S5A**). Staining for androgen receptor showed abundant nuclear staining in most of the luminal epithelial cells in both *Socs1^ΔPE^* and *Socs1^fl/fl^*mice without appreciable difference in staining pattern (**Fig. 2D**). These data indicate SOCS1 deficiency increases the steady-state proliferation of epithelial cells in anterior and dorsal prostate lobes that does not perturb the organization of luminal and basal epithelial cells in glandular acini.

**Figure 2.**
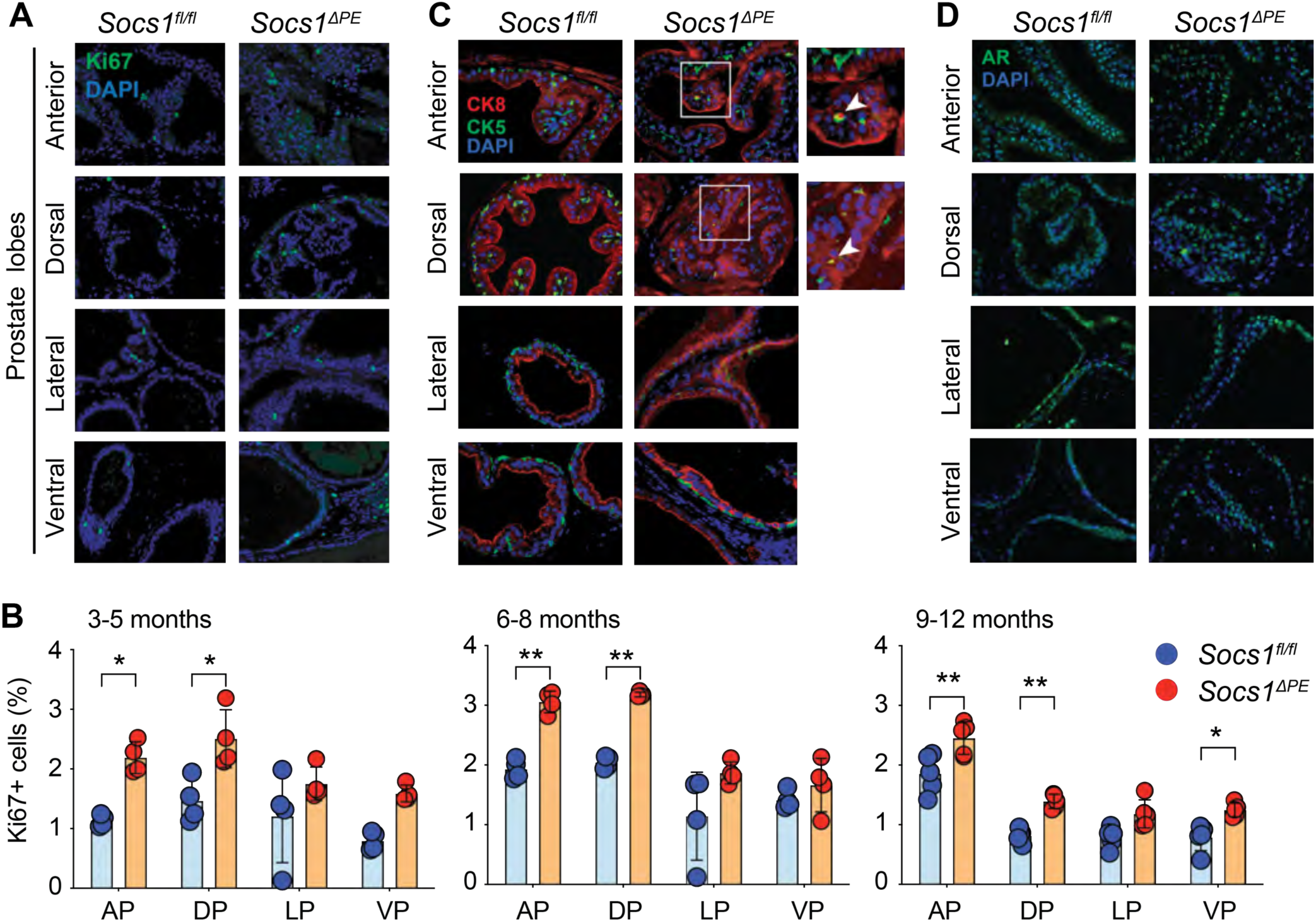
SOCS1 loss increases epithelial cell proliferation without affecting basal and luminal cell distribution. (A,B) Ki67 immunofluorescence staining of prostate gland sections from 6-8 months old *Socs1^fl/fl^* and *Socs1^ΔPE^* mice. (A) Representative regions of anterior, dorsal, lateral and ventral prostate lobes from 3-4 *Socs1^fl/fl^* and *Socs1^ΔPE^* mice. Ki67 staining of prostate glands from 3-5 and 9-12 months-old mice are shown in Supplementary Figure S4. (B) Cumulative data on the proportion of Ki67+ cells in the prostate lobes of different age groups of mice. The number of Ki67+ cells over total number of cells from ten random fields from 4-5 mice per group are shown. Statistics: Mean +SD. Two-way ANOVA with Tukey’s multiple comparison. *p* values: * <0.05; ** <0.01. (C) Immunofluorescence staining of luminal (CK8) and basal (CK5) epithelial cell markers. Representative staining patterns in the prostate lobes of 6-8 months-old mice are shown. CK8 and CK5 staining of 3-5 and 9-12 months are shown in Supplementary Figure S5. Areas of anterior and dorsal lobes of *Socs1^ΔPE^* mice containing CK8+CK5+ cells are enlarged to show the double positive bipotent cells. (D) Androgen receptor staining in 6-8 months old *Socs1^fl/fl^*and *Socs1^ΔPE^* mice.

### Prostate organoids from SOCS1-deficient mice recapitulate the hyperplasia phenotype

To determine whether the increased cell proliferation in the prostate gland of *Socs1^ΔPE^* mice arose from epithelial cell-intrinsic growth deregulation or resulted from co-operation with stromal and immune cells in the tissue microenvironment, we established prostate organoid cultures from enriched prostate epithelial cells (**Fig. 3A**, **Supplementary Fig. 6A**). After 8 days culture, control prostate organoids were large with clear central zones, whereas SOCS1-deficient organoids were smaller and resembled piled up cells (**Fig. 3A**). Size quantification revealed that the latter were only half the size of the control organoids (**Fig. 3B**). H&E staining revealed that most control organoids contained a large lumen surrounded by a single layer of cells, whereas SOCS1-deficient organoids were predominantly compact with multilayered epithelium and much smaller lumen (**Fig. 3C,D**). Moreover, SOCS1-deficient cells yielded significantly more organoids than control cells (**Fig. 3D**). Staining for epithelial cell markers showed that most organoids from both *Socs1^ΔPE^* and *Socs1^fl/fl^* mice contained luminal and basal cells in correct orientation as in the glandular acini, but with more numerous basal cells than in the gland (**Fig. 3E**, **Fig. 2C**). Some of the organoids from both *Socs1^ΔPE^* and *Socs1^fl/fl^*mice contained only basal or luminal cells (**Supplementary Fig S6B**). Notably, SOCS1-deficient organoids contained occasional CK5+CK8+ double positive cells (**Fig. 3E**, **Supplementary Fig. S6B**), as seen in the anterior and dorsal prostate lobes (**Fig. 2C**). Staining for Ki67 and EdU revealed significantly higher number of proliferating cells in SOCS1-deficient organoids compared to control organoids (**Fig. 3F,G**; **Supplementary Fig. S6C**). These data indicate that epithelial cell-intrinsic growth dysregulation is a main cause of prostate hyperplasia in *Socs1^ΔPE^*mice.

**Figure 3.**
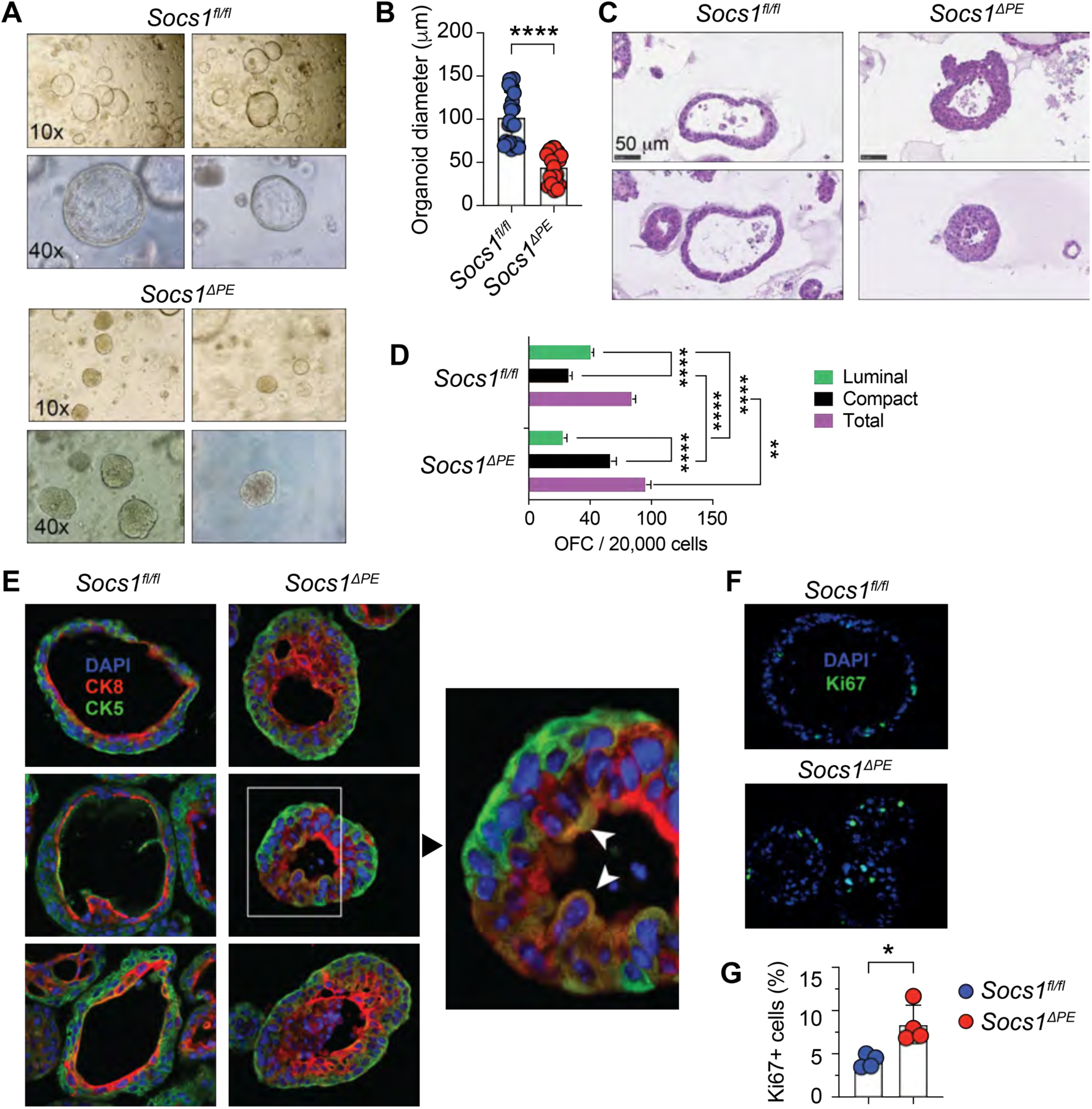
Prostate organoids from SOCS1-deficient mice recapitulate the hyperplasia phenotype. (A) Light microscope images of prostate epithelial organoid cultures established from 5-6 months-old *Socs1^fl/fl^*and *Socs1^ΔPE^* mice on day 8 of culture. Representative cultures from two different mice for each genotype at 10× and 40× magnification are shown. (B) Quantification of organoid size. Twenty-five organoids from 5 mice for each genotype were measured. Mean + SD. Mann-Whitney test. *p* value: * <0.05. (C) Representative images of hematoxylin and eosin-stained sections of FFPE-embedded organoids. (D) Organoid forming count (OFC) per 20,000 prostate epithelial cells seeded in the Matrigel. Morphology of the organoids into luminal and compact type were quantified. Three randomly selected organoid cultures from three mice for each genotype were counted. Statistics: Two-way ANOVA with multiple comparisons test. *p* value: ** <0.01; **** <0.0001. (E) Immunofluorescence staining of luminal (CK8) and basal (CK5) epithelial cell markers. Three representative prostate organoids from *Socs1^fl/fl^* and *Socs1^ΔPE^* mice are shown. The boxed area in one of the SOCS1-deficient organoids is enlarged in the right panel to indicate double positive (CK8+CK5+) cells (arrowheads). (F) Representative Ki67 staining of proliferating cells in SOCS1-deficient and control prostate organoids. (G) Quantification of Ki67+ cells. Mean + SD. Unpaired *t* test. *p* value: **** <0.0001.

### SOCS1 expression in prostate epithelial cells regulates COX-2 expression

EGF is a key growth promoter in normal and hyperplastic prostate gland ^35,48^. In addition to providing growth stimulatory signals, EGF receptor activation upregulates cyclooxygenase-2 (COX2) needed for the biosynthesis of prostaglandin E2 (PGE2), which is associated with cell growth in prostate cancer and many other solid tumors, contributing to tumor aggressiveness and resistance to therapy ^49–52^. Hence, we examined COX-2 expression in the prostate glands of *Socs1^fl/fl^* and *Socs1^ΔPE^* mice. Whereas COX-2 expression was non-detectable in the prostate glands of four months-old *Socs1^fl/fl^* mice, luminal epithelial cells in all four prostate lobes of *Socs1^ΔPE^* mice of the same age showed marked expression of COX-2 (**Fig. 4A**). Increased COX-2 expression in SOCS1-deficient prostate glands persisted through twelve months of age, when only a few dorsal and ventral lobes of prostates from control mice showed discernible COX2 staining (**Fig. 4B**). These results indicate that SOCS1 expression inhibits a key mitogenic and potentially pro-oncogenic signaling pathway in prostate epithelial cells.

**Figure 4.**
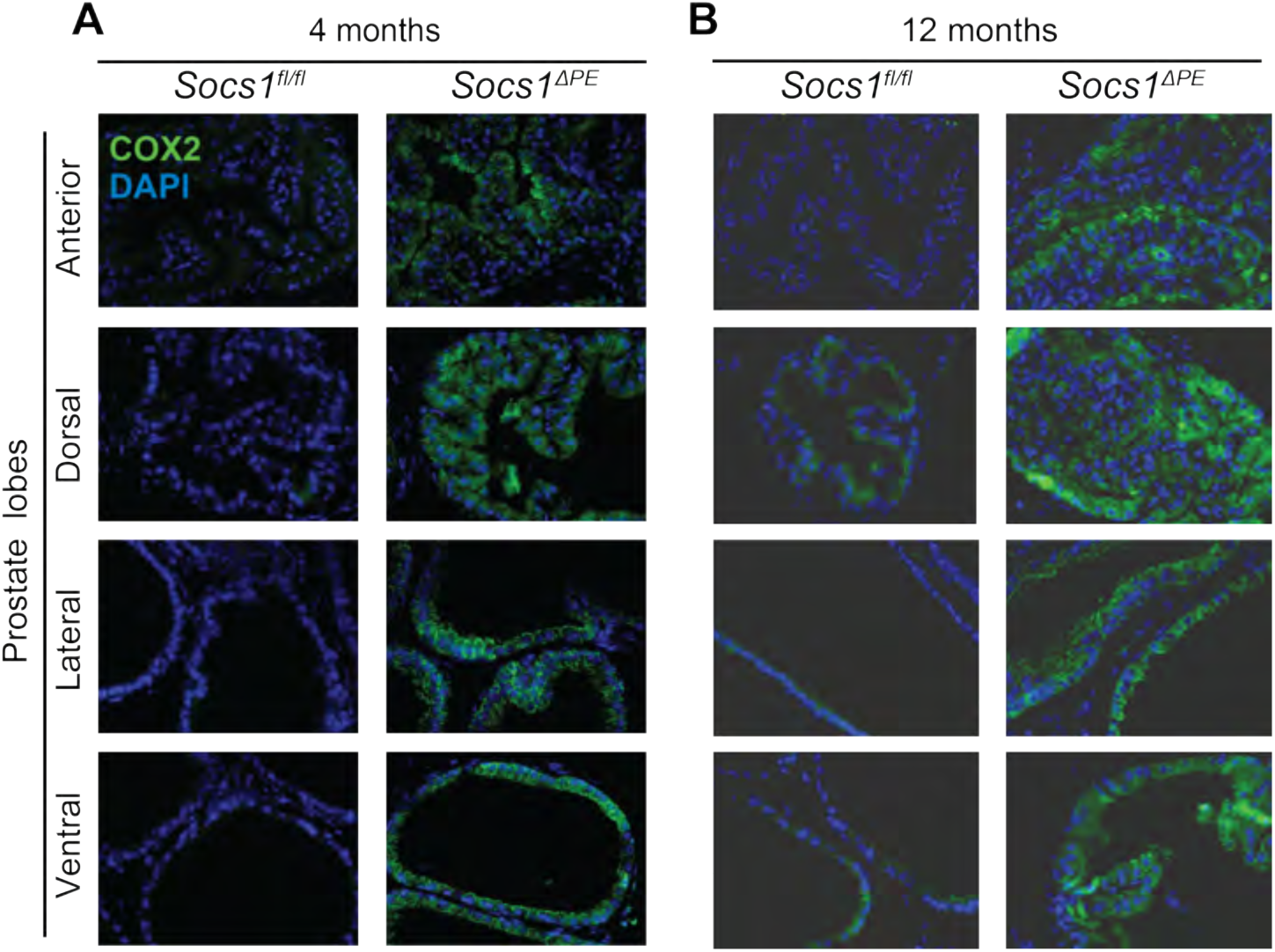
Upregulation of COX-2 expression in SOCS1-deficient prostate glands. Immunofluorescent staining of COX-2 in FFPE-sections of prostate glands from 4 (A) and 12 (B) months-old *Socs1^fl/fl^* and *Socs1^ΔPE^* mice. Representative regions of anterior, dorsal, lateral and ventral prostate lobes from 3-4 mice per group are shown.

### High-fat diet induces pre-neoplastic changes in SOCS1-deficient prostate gland

Population studies suggest that consumption of high-fat diet (HFD) carries a risk for aggressive prostate cancer through promoting obesity and systemic inflammation ^53^. Many studies on cellular, transgenic and knockout mouse models have shown that high-fat diet elicits inflammation, oxidative stress and production of cytokines, chemokines and growth factors, promoting proliferation of epithelial and stromal cells that contribute to prostate cancer progression ^43,54–57^. To test whether HFD could synergize with SOCS1-deficiency and lead to neoplasia, we fed *Socs1^ΔPE^* and control mice with HFD (>60% energy from fat) *ad libitum* or normal control diet (CD), from 6 weeks of age and examined the prostate glands ten months later ^43^. Both groups of HFD-fed mice showed comparable body weight gain (**Supplementary Fig. S7A**) and did not show gross morphological abnormality of the prostate gland. However, more than half of HFD-fed *Socs1^fl/fl^* mice displayed appreciable changes in prostate histology featuring hyperplasia and thickening of the fibromuscular layer in anterior and dorsal lobes, and inflammatory cell infiltration and fibrotic changes in lateral and ventral prostate lobes compared to CD-fed controls (**Supplementary Fig. S7B-E**). HFD exacerbated hyperplasia in *Socs1^ΔPE^* mice, characterized by irregular dysplastic nuclei in anterior and dorsal lobes resembling PIN that did not affect the lobular architecture (**Supplementary Fig. S7B-E**). Such nuclear abnormality was rarely seen in HFD-fed *Socs1^fl/fl^* mice. These data indicate that SOCS1 prevents the development of pre-neoplastic lesions under conditions of heightened inflammation caused by HFD.

### SOCS1 loss in prostate epithelial cells increases susceptibility to cancer development

Despite increasing cell proliferation and susceptibility to undergo pre-neoplastic changes in an inflammatory setting caused by HFD, SOCS1 deficiency in prostate epithelial does not elicit neoplasia. To investigate whether SOCS1 deficiency in prostate epithelial cells could synergize with other potentially oncogenic stimuli, we infected 3-4 months-old *Socs1^fl/fl^* and *Socs1^ΔPE^*mice with uropathogenic *Escherichia coli* UPEC1677, which harbors the *pks* island containing *clb* genes coding for the biosynthesis of colibactin ^58,59^. Colibactin is a genotoxin known to cause DNA double strand breaks and promote carcinogenesis in mouse models of colorectal cancer, and induce TMPRSS2-ERG fusion oncoprotein in prostate cancer cell lines ^60–62^. Prostate glands from most of the UPEC1677-infected *Socs1^ΔPE^* mice presented large tissue masses without recognizable lobes, whereas *Socs1^fl/fl^*mice did not present any change in gross morphology (**Fig. 5A**). However, H&E-stained tissue sections of the UPEC1677-infected *Socs1^fl/fl^* mice revealed marked hyperplasia and dysplasia in all four prostate lobes, and intraepithelial neoplasia in anterior and dorsal lobes (**Fig. 5B,C**). On the other hand, the glandular architecture was completely obliterated in SOCS1-deficient prostate replaced by well recognizable tumor nodules, indicating invasive cancer (**Fig. 5B,C**). Such changes were observed in 19 out of 21 *Socs1^ΔPE^* mice, whereas only 1 out of 14 *Socs1^fl/fl^*mice developed cancer (**Fig. 5D**). In vivo labelling of infected mice with EdU to identify S phase cells incorporating the DNA analog, and combined Ki67 and EdU staining of tumor sections revealed numerous Ki67+Edu-, Ki67-Edu+ and Ki67+Edu+ cells undergoing proliferation within the prostate tumor nodules of *Socs1^ΔPE^* mice whereas the dysplastic prostates of *Socs1^fl/fl^* mice contained fewer proliferating cells (**Supplementary Fig. S8**). Whereas the distribution of CK8+ luminal and CK5+ basal epithelial cells in the prostate glands of *Socs1^fl/fl^* mice remained organized in glandular acini, the tumor nodules of *Socs1^ΔPE^* mice displayed complete disorganization of these cells with readily detectable CK8+CD5+ cells, and some nodules containing large pockets of double positive cells (**Fig. 5E**). Marking the stromal fibroblasts in the fibromuscular layer surrounding the glandular acini by staining the tissue sections for αSMA revealed discontinuous and fragmented staining in the prostate tissues of *Socs1^ΔPE^* mice, indicating extensive breaching of the basement membrane, which were intact in the prostates of *Socs1^fl/fl^* mice (**Fig. 5F**). These results indicate that SOCS1 deficiency in prostate epithelial cells can synergize with potentially oncogenic stimuli such as the ones arising from genotoxic stress to initiate neoplasia and its progression towards invasive cancer.

**Figure 5.**
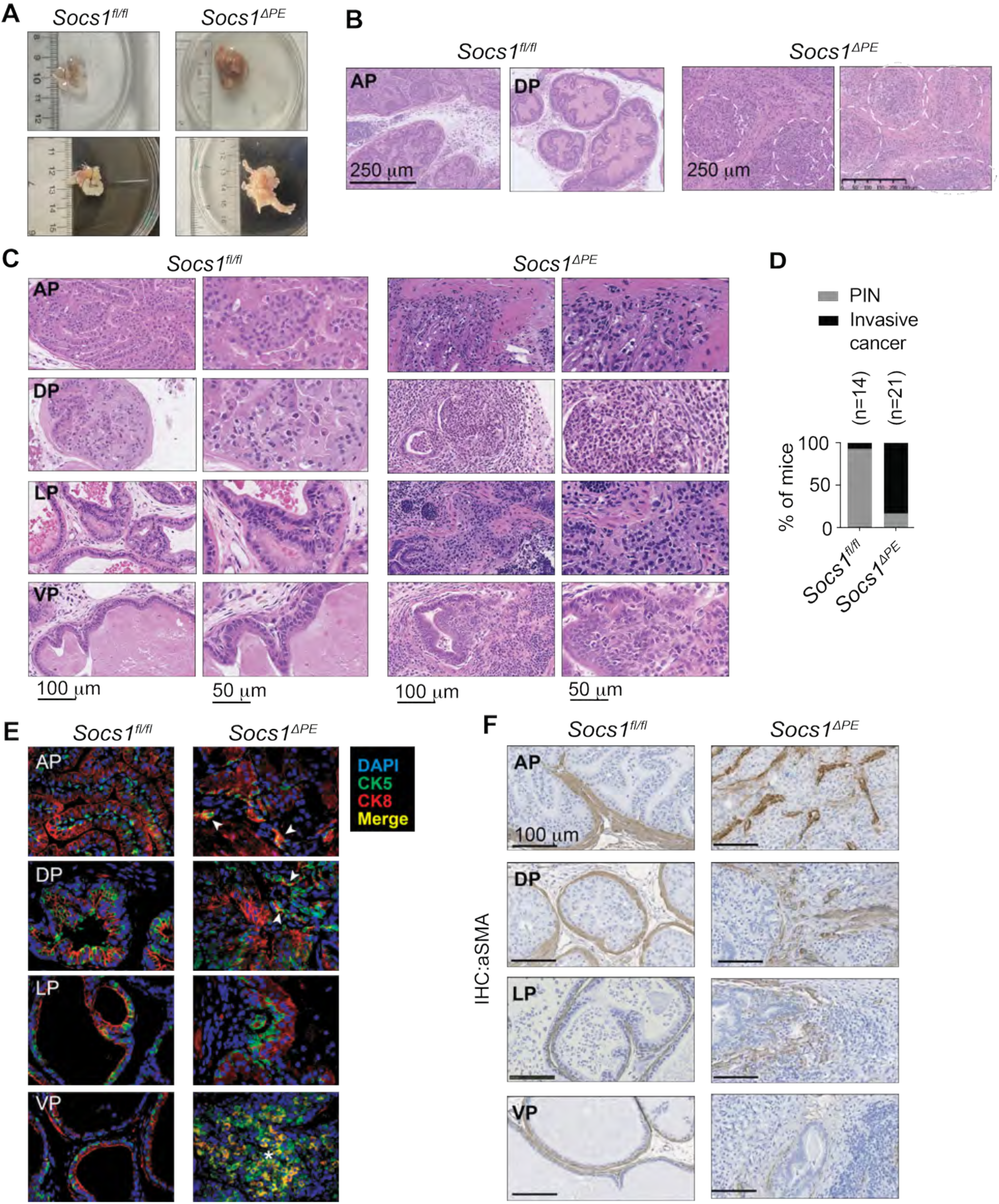
*Socs1^ΔPE^*mice infected with a genotoxic bacteria develop prostate cancer. *Socs1^fl/fl^*and *Socs1^ΔPE^* mice were infected at 3-4 months of age with uropathogenic *E. coli* UPEC1677 via transurethral route. When some of the infected animals showed signs of discomfort and malaise, around 6-7 months after infection, all mice were euthanized and their prostate glands were examined. (A) Dissected prostate tissues from two representative mice for each genotype. Seminal vesicles are not removed from prostates of control *Socs1^fl/fl^* mice. The boundary of the prostate gland in *Socs1^fl/fl^* mice is indicated by dotted lines. (B) FFPE sections of prostate tissues were stained by hematoxylin and eosin. Low magnification images showing tumor nodules in *Socs1^ΔPE^* mice and intact prostate lobular architecture in *Socs1^fl/fl^* mice. (C) Higher magnification images of the prostate lobes showing extensive hyperplasia, dysplasia and intraepithelial neoplasia in the prostate lobes of *Socs1^fl/fl^* mice and disorganized tumor nodules with extensive extracellular matrix deposition in the neoplastic prostate tissues *Socs1^ΔPE^* mice. (D) Cumulative data on cancer incidence in *Socs1^ΔPE^* mice and prostate intraepithelial neoplasia (PIN) in *Socs1^fl/fl^* mice. (E) Immunofluorescence staining of CK8+ luminal and CK5+ basal epithelial cells. Occasional double positive cells in the tumor nodules of *Socs1^ΔPE^* mice are indicated by arrowheads, and the asterisk indicates pockets of CK8+CK5+ cells in some tumor nodules. (F) Immunohistochemical staining of alpha smooth muscle actin (αSMA) in prostate tissue sections. Representative data from 3-4 mice per group.

### SOCS1 deficiency in prostate epithelial cells increases DNA damage induced by colibactin

The heightened susceptibility of *Socs1^ΔPE^* mice to develop prostate cancer following colibactin producing UPEC1677 suggested that the loss of SOCS1, which is implicated in p53 activation ^63^, may impact DNA damage repair pathways following colibactin-induced DNA damage. To test this hypothesis, we deleted the *clbP* gene involved in a key step in colibactin biosynthetic pathway ^64^ and confirmed the inability of UPEC1677*ΔclbP* to cause DNA damage in the BPH-1 human benign prostatic hyperplasia cell line by γH2AX staining (**Supplementary Fig. S2, S9**). Next, we compared the ability of UPEC1677 and UPEC1677*ΔclbP* to cause DNA damage in epithelial cell monolayers derived from *Socs1^fl/fl^* and *Socs1^ΔPE^*mouse prostate organoids. UPEC1677 but not UPEC1677*ΔclbP* caused significant DNA damage in *Socs1^fl/fl^* epithelial cells, which was significantly increased in *Socs1^ΔPE^* cells (**Fig. 6A-B**). Transurethral prostate infection with UPEC1677*ΔclbP* failed to induce any malaise or discomfort up to 8 months post-infection, and the prostate glands of these mice did not show gross morphological abnormalities (**Fig. 6C**). However, histological examination of the prostate glands revealed extensive immune cell infiltration across all prostate lobes in both *Socs1^fl/fl^* and *Socs1^ΔPE^* mice (**Fig. 6D-E; Supplementary Fig. S10**). Anterior and dorsal lobes showed extensive epithelial cell remodeling, glandular distortion and pre-neoplastic changes in both *Socs1^fl/fl^*and *Socs1^ΔPE^* mice (**Fig. 6E**). These data indicate that SOCS1 in prostate epithelial cells attenuates colibactin-induced DNA damage and thereby confers resistance to carcinogenesis induced by genotoxic agents.

**Figure 6.**
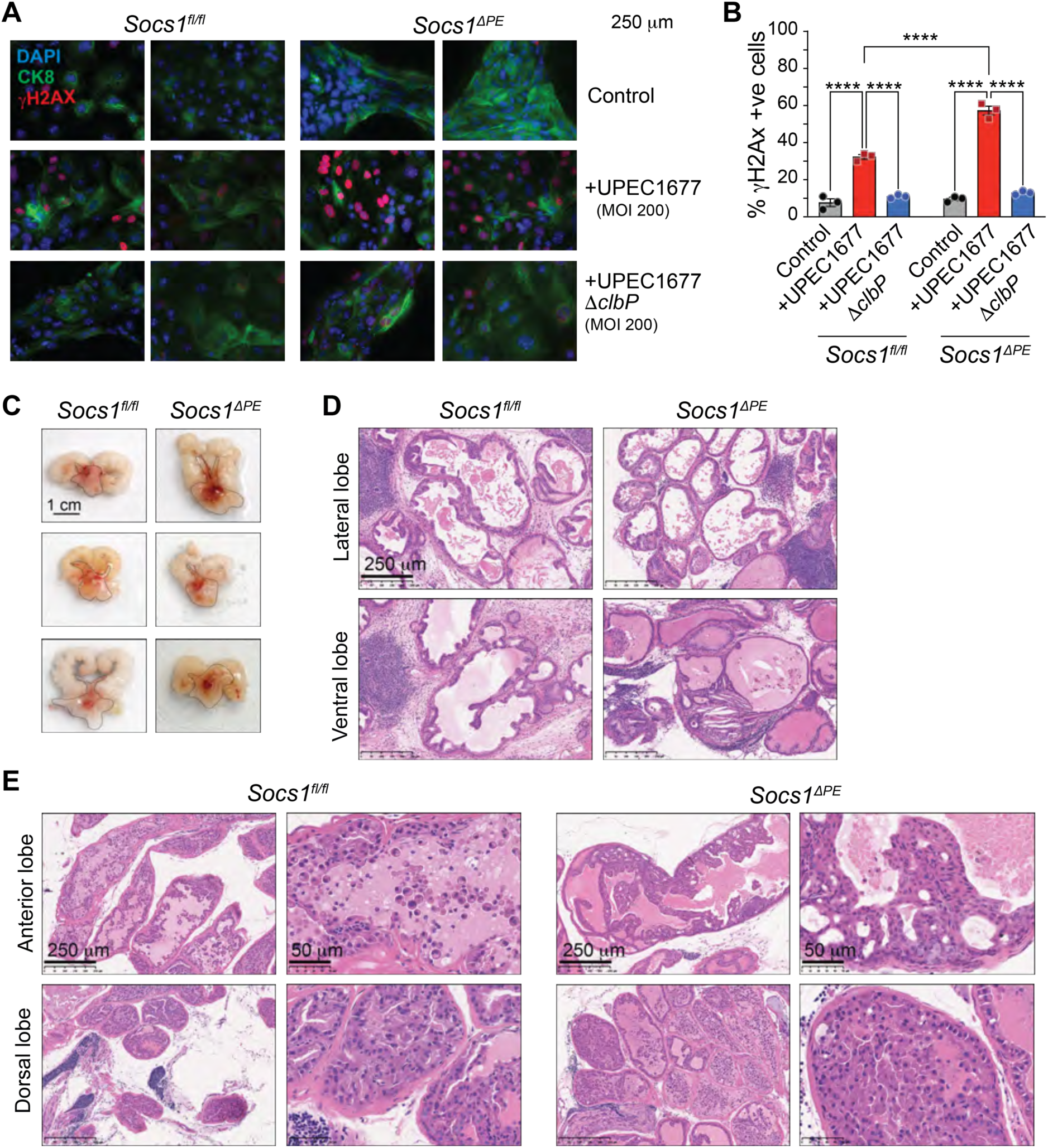
SOCS1 deficiency in prostate epithelial cells increases DNA damage induced by colibactin. (A) Prostate organoids from *Socs1^fl/fl^* and *Socs1^ΔPE^*mice were disrupted and grown as epithelial cell monolayers grown on glass coverslips. The cells were exposed to UPEC1677, or UPEC1677*ΔclbP* defective in colibactin biosynthesis, for 30 min, washed and cultured for additional 18 h before fixation and immunofluorescence staining for γH2AX and CK8. After staining the nuclei with DAPI the cells were examined under a fluorescence microscope. Two representative images for each condition from two separate experiments are shown. (B) quantification of γH2AX positive cells over total DAPI-stained nuclei. Cumulative data from three separate experiments are shown. Mean + SEM. Two-way ANOVA with multiple comparisons test. *p* value: **** <0.0001. (C) Prostates of 4 month-old *Socs1^fl/fl^* and *Socs1^ΔPE^* mice (n=4-6) were inoculated with UPEC1677*ΔclbP* and their prostate glands were examined 8 months later. Images of the entire urogenital apparatus including seminal vesicles from three mice in each genotype are shown, with the boundary of the prostate gland delineated by dotted lines. (D,E) Hematoxylin and eosin-stained sections of lateral and ventral prostate lobes (D), and anterior and dorsal prostate lobes (E) from UPEC1677*ΔclbP*-infected *Socs1^fl/fl^*and *Socs1^ΔPE^* mice. Representative images from 4 mice for each genotype are shown (scale bar 250 μm). Higher magnification images are shown for anterior and dorsal lobes in (E) (scale bar 50 μm).

### SOCS1 loss dysregulates epithelial lineage markers and pro-neoplastic signaling proteins

To gain insight into molecular pathways dysregulated by SOCS1 deficiency in prostate epithelial cells, we performed proteomic analysis on prostate organoids established from *Socs1^fl/fl^* and *Socs1^ΔPE^*mice. Quality control analysis on quadruplicate samples using PCA plot, sample-to-sample correlation and Euclidean heatmaps revealed clear internal grouping of SOCS1-deficient and control cells, with more than 7100 proteins detected in each sample (**Supplementary Fig. S11**). Volcano plot analysis showed 219 upregulated and 47 downregulated proteins in SOCS1-deficient cells compared to control cells with an FDR value of <0.1 that included 79 upregulated and 13 downregulated proteins at FDR 0.05 (**Fig. 7A**). Notably semenogelin 1 (SEMG1), seminal vesicle secretory protein 6 (SVS6) and SVS3A, which are primarily expressed in seminal vesicles, but can also be expressed in prostate cancer ^65,66^, are downregulated in SOCS1-deficient cells (**Fig. 7A**). Analysis of prostate epithelial lineage markers, following the published gene signatures of basal epithelial cells and luminal epithelial cells of anterior, dorsolateral and ventral prostate lobes ^67,68^, revealed numerous upregulated and downregulated proteins in SOCS1-deficient organoids compared to control organoids (**Supplementary Figure S12**). Notable among them are downregulation of KRT7 and KRT15, and upregulation of NECTIN1, SPARC, CAVIN3, LCN2, and SERPINH1 for basal cell markers, and downregulation of PBSN and PDLIM2, and upregulation of TGM4, SARAF and PROM2 for luminal cell markers, among several others ^67^. However, one SOCS1-deficient organoids (KO4) not only resembled all other replicates of this group for upregulated proteins but also showed similarity to the upregulated proteins of control organoids, which is also reflected its partial overlap with control samples in the PCA plot (**Supplementary Fig. S10, S12**). A gene set enrichment analysis (GSEA) of cancer hallmark pathways revealed positive enrichment of Interferon Gamma Response, Inflammatory Response, Il2-Stat5 Signaling and Pi3k Akt Mtor Signaling, and negative enrichment of P53 pathway, Dna Repair and Xenobiotic Metabolism, among the significantly modulated proteins in SOCS1-deficient cells (**Fig. 7B, Supplementary Figures S13 and S14**), commensurate with the known regulatory functions of SOCS1 ^24,25,69,70^. Even though enrichment of these pathway gene sets were not statistically significant (FDR >0.05), proteins modulated by SOCS1 deficiency in these pathways could potentially contribute to the hyperplastic phenotype and sensitivity to colibactin-induced DNA damage of SOCS1-deficient cells (**Fig. 3, 6**). GSEA using KEGG revealed significant negative enrichment of ‘Rna Polymerase’ and ‘Pyrimidine metabolism’ pathways (**Fig. 7C**). However, heatmap analysis of proteins significantly modulated by SOCS1 within the pyrimidine metabolism pathway revealed upregulation of proteins that mediate de novo synthesis (CAD, CTPS2, DHODH) and salvage pathway (NT5C1A, NME4, CDA, NT5E, ENTPD5, NME3), RNA polymerase II isoforms (POLR3A, POLR3D, POLR3E), DNA polymerase isoforms (POLE2, POLE3), which are associated with rapid cell growth and proliferation in cancer (**Fig. 7D**). Furthermore, SOCS1-deficient cells showed a positive enrichment of Prostate Cancer and Pathways in cancer gene sets in SOCS1-deficient cells, which, though not significant (FDR >0.05) included upregulation of PI3K/AKT/mTOR Pathway (IGF1R, PTEN, AKT2, MTOR), MAPK/ERK Pathway (ERBB2, ARAF, RAF1, MAPK3) and Wnt/b-catenin Pathway (CTNNB1, CHUK) proteins, which could contribute to increased cell survival, growth and proliferation (**Fig. 7E**).

**Figure 7.**
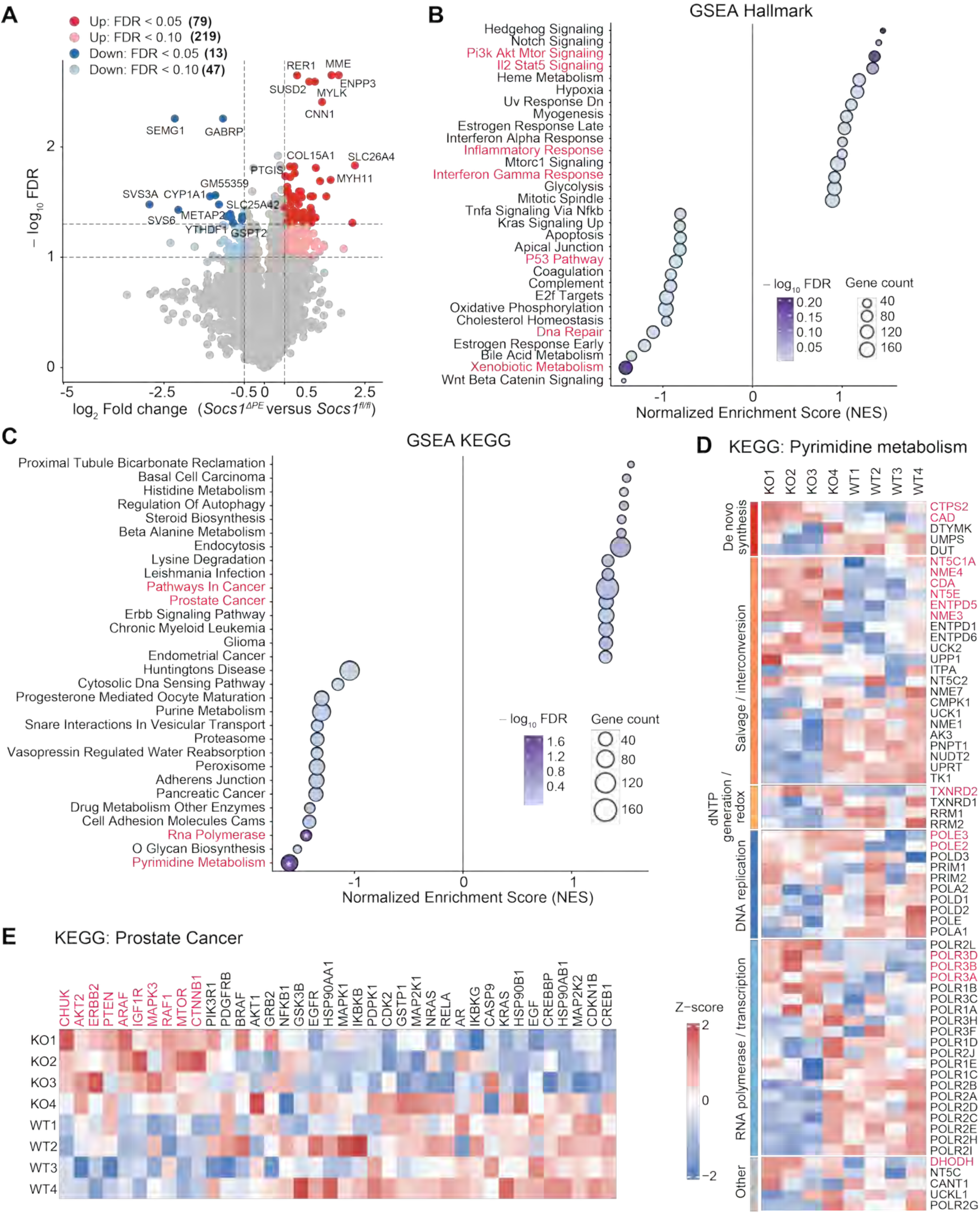
Signaling pathways modulated in SOCS1-deficiency prostate epithelial organoids. Total proteome analysis was performed on prostate epithelial cells from organoids established from *Socs1^fl/fl^*mice (n=4) and *Socs1^ΔPE^* mice (n=4) collected on day 8 of culture. PCA plots and quality control data are shown in Supplementary Figure S10. (A) Volcano plot showing differentially expressed proteins (DEPs) in SOCS1-deficient cells compared to control cells. DEPs showing fold change (FC) ≤-1 and ≥1 and adjusted *p*-value (False discovery rate, FDR) <0.05 and <0.1 are color-coded. (B) Enrichment of GSEA hallmark pathways proteins among DEPs modulated in SOCS1-deficient cells. (C) Enrichment of GSEA KEGG pathways proteins among DEPs modulated in SOCS1-deficient cells. Asterisks indicate enriched pathways with FDR <0.05. (D) Heatmap representation of DEPs within the functional modules of KEGG Pyrimidine Metabolism pathway. KO, cells from *Socs1^ΔPE^* mice; WT, cells from *Socs1^fl/fl^* mice. (E) Heatmap of DEPs within the KEGG Prostate Cancer pathway.

## Discussion

SOCS1 exerts diverse cellular functions that can guard against the development and progression of cancer. These functions include inhibition of inflammatory cytokine production, attenuation of oncogenic growth factor signaling, regulating oxidative stress response and facilitating p53 activation. In the present study, we show using the prostate-specific SOCS1 knockout mouse model that SOCS1 expression in prostate epithelial cells is essential to preserve post-pubertal tissue homeostasis, restrain age-associated hyperplasia development and confer protection against neoplastic transformation induced by genotoxic stress.

Earlier studies on SOCS1-deficient mice have demonstrated the tumor suppressor role of SOCS1 in liver hepatocellular carcinoma and colorectal cancer induced by chemical carcinogens ^71–73^. In the TCGA transcriptomic data, higher SOCS1 expression predicts better survival probability in liver cancer and certain other cancers, whereas no such correlation was observed for several other cancers such as colon or rectal adenocarcinoma and prostate cancer. The correlation is even negative for a few other cancers. This conundrum could arise from the induction *SOCS1* gene expression in diverse cell types within the tumor microenvironment by various stimuli, and inhibition of SOCS1 protein expression by microRNAs and SOCS1 functions by post-translational modifications ^28,29,74^. We have shown earlier that SOCS1 protein expression in prostate cancer specimens inversely correlates with increasing tumor grade and metastasis even though SOCS1 gene expression in TCGA of prostate adenocarcinoma does not correlate with the survival probability ^31^. Nonetheless, the current genetic study demonstrates the tumor suppressor role of SOCS1 that likely acts at the level of both tumor initiation and its progression. While SOCS1 gene loss alone is not sufficient to cause neoplasia, it can synergize with another pro-oncogenic signal such as the one elicited by the bacterial genotoxin colibactin.

A key finding of our study is the progressive hyperplastic phenotype caused by SOCS1 deficiency in the adult prostate without affecting early prostate development, maturation or growth. The Pb-Cre4 transgene is expressed in prostate epithelial cells within one week after birth and is active throughout sexual maturation and adult life, with elevated expression in lateral and ventral lobes compared to anterior and dorsal lobes, and higher expression in luminal cells (∼85%) than in basal cells (∼50%) ^33,75,76^. Because *Socs1^ΔPE^* mice did not display any discernible histological changes in the prostate lobes until after 3 months of age well after weaning and attaining puberty, and did not display any notable alterations in the distribution of luminal and basal cells, the progressive hyperplasia phenotype could arise from at least two mutually non-exclusive factors. Firstly, as SOCS1 is a key regulator of cytokine and growth factor signaling, loss of SOCS1 could increase the responsiveness of epithelial cells to androgen-induced stromal cell-derived growth factors during tissue remodelling of the post-pubertal prostate gland ^41,77^. Circumstantial evidence supporting this possibility include induction of SOCS1 by androgens, development of PIN in the HGF receptor MET transgenic mice and attenuation of MET signaling by SOCS1 ^26,45,69^. However, the incidence of hyperplasia occurring randomly over a period of nine months suggests a second contributing factor, possibly exposome stimuli arising from environmental exposure. Even though *Socs1^ΔPE^* mice were housed in specific pathogen-free conditions, progressive establishment of the gut microbiota after weaning ^78^, and possibly that of the prostate microbiota, could increase systemic and local availability of microbial products such as bacterial lipopolysaccharide (LPS). SOCS1 inhibits LPS signaling via TLR4 and attenuates production of inflammatory and growth promoting cytokines such as TNFα and IL-6 ^22,23^. Prostate epithelial cells express TLR4 and respond to LPS ^79^. IL-6 and TNFα are implicated in promoting prostate epithelial cell proliferation, and transgenic expression of IL-6 in the mouse prostate gland induces spontaneous cancer development ^18,19^. Intestinal microbiota, which is implicated in prostate cancer progression ^80^, may also contribute to the hyperplasia phenotype in *Socs1^ΔPE^* mice through gut-derived microbial metabolites reaching the prostate tissue via systemic circulation.

Prostate glands of *Socs1^ΔPE^* mice display hyperplasia predominantly in anterior and dorsal lobes with negligible inflammatory cell infiltration, whereas the latter is more prevalent in ventral and lateral lobes. Moreover, hyperplastic changes in the prostate begin earlier than the inflammatory cell infiltration in *Socs1^ΔPE^* mice. Such spatial and temporal segregation of hyperplasia and inflammation suggests that signaling events dysregulated by SOCS1 deficiency may differ in different lobes, possibly influenced by differential stromal cell content and cellular heterogeneity ^81,82^. The prostate organoid data suggest that the hyperplastic phenotype of SOCS1-deficient prostate is driven largely by epithelial cell-intrinsic mechanisms. SOCS1-deficient epithelial cells generate more organoids, show increased cell proliferation, and form mainly compact, multilayered structures instead of the larger hollow organoids formed by control cells. Mouse prostate organoids are heterogeneous and reflect the epithelial state of the cells that generate them. Luminal progenitors alone can efficiently generate organoids, often with hollow lumens containing both luminal and basal epithelial cells, while other epithelial states carrying pro-neoplastic genetic lesions can produce more solid or irregular structures ^83,84^. In this context, the compact solid phenotype of SOCS1-deficient prostate epithelial organoids likely results not only from increased proliferation but also from a shift in epithelial identity or progenitor composition. This notion is supported by more frequent occurrence of CK8+CK5+ cells in SOCS1-deficient prostate epithelial organoid cultures and dysregulated expression of basal and luminal epithelial cell markers in the proteome analysis. CK8+CK5+ cells are viewed as intermediate or bipotent progenitor-like epithelial cells and have been described during prostate development, differentiation, and epithelial regeneration ^85,86^. Rare transitional epithelial states have also been detected by lineage-tracing approaches in the adult prostate ^87^. Because the Pb-Cre driver mainly targets luminal epithelial cells ^33,75,76^, the increase in double-positive cells after *Socs1* deletion raises the possibility that SOCS1-deficient luminal-lineage cells may acquire basal features or enter an intermediate state. This interpretation is supported by studies showing that inflammation and tissue damage can promote epithelial plasticity, including expansion of double-positive transit-amplifying cells and enhanced basal-to-luminal differentiation ^88^. Thus, SOCS1 may help maintain luminal lineage conformity in the adult prostate and its loss may create a permissive state for epithelial plasticity. Overall, the progressive increase in epithelial tufting and multilayering in the post-pubertal prostates of *Socs1^ΔPE^*mice supports a model in which SOCS1 acts as a regulator of both cell proliferation and differentiation for long-term maintenance of epithelial equilibrium rather than as a developmental determinant. Independent of this function, SOCS1 serves to reduce chronic inflammatory stress in the prostate, although mechanisms underlying the spatial segregation of SOCS1 functions remain to be clarified.

SOCS1 deficient prostates do not develop spontaneous neoplasia despite showing increased epithelial cell proliferation, dysregulated plasticity, architectural disruption and elevated COX-2 expression. COX-2 expression and prostaglandin signaling have been linked to epithelial proliferation, inflammation and carcinogenesis in the prostate and other solid tumors ^49–52^. Moreover, SOCS1 deficiency appears to create a pre-neoplastic state that is permissive to transformation. Proteins involved in PI3K/AKT/mTOR Pathway (IGF1R, PTEN, AKT2, MTOR), MAPK/ERK Pathway (ERBB2, ARAF, RAF1, MAPK3) and Wnt/b-catenin Pathway (CTNNB1, CHUK), which are implicated in cell survival, growth and proliferation, are enriched in the proteome SOCS1-deficient cells. Furthermore, SOCS1-deficient cells show increased expression of proteins involved in de novo (CTPS2, CAD) and salvage (NT5C1A, NT5E, NME3, NME4, CDA, ENTPD5, ENTPD6, UCK2) pathways of pyrimidine metabolism, DNA replication (POLE3, POLE2, POLD3) and DHODH, all of which are implicated in prostate cancer progression ^89,90^. Higher steady state activation of these pathways in SOCS1-deficient cells may underlie the development of invasive prostate cancer following transurethral infection of *Socs1^ΔPE^* mice with colibactin-producing UPEC1677, while control prostates developed hyperplasia, dysplasia or PIN-like changes. Colibactin is a genotoxin that induces DNA double-strand breaks and promotes colorectal cancer and induces oncogenic TMPRSS2:ERG fusion protein in prostate cancer cell lines ^60–62^. Our finding that UPEC1677 mutant incapable of producing colibactin failed to cause cancer in *Socs1^ΔPE^*mice, and that SOCS1-deficient prostate organoid-derived epithelial cells accumulated more DNA damage upon exposure to colibactin-producing bacteria, strongly support the idea that SOCS1 protects prostate epithelial cells from genotoxic injury. In this context, SOCS1 has been linked to p53 activation and stress responses ^63,73^, and proteins involved in DNA repair, p53 pathway and xenobiotic metabolism are dysregulated in SOCS1-deficient cells with some upregulated and several downregulated proteins. Overall, SOCS1 expression in prostate epithelial cells could confer protection against carcinogenesis by attenuating potentially oncogenic signaling pathways while promoting anti-neoplastic cellular pathways.

In summary, our findings support a model in which SOCS1 maintains prostate epithelial homeostasis through multiple complementary mechanisms. SOCS1 attenuates reduces survival and growth-promoting cytokine and growth factor signaling, limits chronic epithelial cell proliferation, preserves luminal cell identity, prevents the emergence of intermediate progenitor-like states, and protects against DNA damage–driven malignant transformation. Loss of functional SOCS1 in prostate epithelial cells through gene repression, mutations or functional inactivation by post-translational modifications could compromise these functions leading to increased cell proliferation, disruption of homeostasis, inflammation and increased susceptibility to carcinogenesis following genotoxic challenge. Furthermore, the *Socs1^ΔPE^* mouse model provides a valuable platform to investigate mechanisms of prostate hyperplasia and initiation and progression of cancer by microbial genotoxins.

## Supporting information

Protein expression data_All replicates

Ranked Lineage markers

## Ethics statement

All experiments on mice were carried out with the approval of the Université de Sherbrooke Ethics Committee for Animal Care and Use (Protocol ID: 2020-2502 and 2024-4410).

## Acknowledgements

AUH received Bourse d’excellence VoiceAge, and MN and MM received Bourse Abdenour-Nabid doctoral scholarships from the Faculty of Medicine and Health Sciences, Université de Sherbrooke. The authors thank Mr. Dominique Lévesque of the Université de Sherbrooke Mass spectrometry platform for help with proteomics data processing.

## Authors’ Contributions

Conceptualization, S.I., G.F., S.R., and A.M.; Methodology, A.U.I., M.N., A.A.C., E.N.K.L., A.M., and S.R.; formal analysis, A.U.I., M.N., M.M., D.B.J., S.P., S.R. and S.I.; resources, A.Y., E.M., A.M., S.R. and S.I.; data curation, A.U.I., M.N., M.M., and S.I.; writing—original draft preparation, A.U.I., M.N., M.M., and S.I.; writing—review and editing, A.U.I., M.N., M.M., E.N.K.L., D.B.J., S.P., A,Y., G.F., A.M., S.R., and S.I.; supervision, E.M., S.R. and S.I.; funding acquisition, G.F., A.M., and S.I.

## Data Availability

Curated mass spectrometry data of intensity profiles of proteins detected in all biological replicates, and expression levels of epithelial lineage markers are given in supplementary spreadsheets. Materials used in this research are available upon request.

## Abbreviations

BPH: benign prostate hyperplasia
CFU: colony forming units
FFPE: Formalin-fixed paraffin-embedded
LPS: lipopolysaccharide
MOI: multiplicity of infection
NDP: Nanozoomer Digital Pathology
OFC: organoid formation count
PFA: paraformaldehyde
PIA: proliferative inflammatory atrophy
PIN: prostate intraepithelial neoplasia
αSMA: alpha smooth muscle actin
SOCS1: suppressor of cytokine signaling 1
UPEC: uropathogenic *Escherichia coli*.

