## Supplementary Tables for "SOCS1 expression in prostate epithelial cells is essential for tissue homeostasis and tumor suppression"

Ihsan et al.,

**Supplementary Table S1.** Antibodies used for immunofluorescence and immunohistochemistry.

| 1° Antibody | Species | Source | Dilution |
|---|---|---|---|
| CK5 | Rabbit pAb | Biolegend # 905504 | 1:200 |
| CK8 | Mouse mAb | Biolegend # 904804 | 1:100 |
| γ-H2AX | Rabbit polyclonal | Abcam #AB11174 | 1:500 |
| Ki67 | Rabbit mAb | Cell Signaling # 12202 | 1:600 |
| αSMA | Mouse mAb | Abcam # 7817 | 1:500 |

| 2° Antibody | Source | Dilution |
|---|---|---|
| Alexa Fluor 488 goat Anti-rabbit IgG (H+L) | *ThermoFisher* # A-11034 | 1:500 |
| Alexa Fluor 488 goat Anti-rat IgG (H+L) | *ThermoFisher* # A-11006 | 1:500 |
| Alexa Fluor 488 goat Anti-mouse IgG (H+L) | *ThermoFisher* # A-11029 | 1:500 |
| Alexa Fluor 568 goat Anti-mouse IgG (H+L) | *ThermoFisher* # A-11031 | 1:500 |
| Alexa Fluor 568 goat Anti-rabbit IgG (H+L) | *ThermoFisher* # A-11036 | 1:500 |

**Supplementary Table S2.**
Primers used for the construction of the UPEC1677Δ*clbP* strain.

| Primer | Description | Sequence |
|---|---|---|
| EM5550 | Forward for deleting *clbP* | GGTGTTCACAGGATGACAATAATGGAACACGTTAGCATTAGTGTAGGCTGGAGCTGCTTC |
| EM5551 | Reverse for deleting *clbP* | AAACAATACAAACTGATATTACTCATCGTCCCACTCCTTGCATATGAATATCCTCCTTAG |
| EM5554 | Forward for sequencing *clbP* | GTCAACAAGGCAAGCGGCTA |
| EM5555 | Reverse for sequencing *clbP* | CCTTTCTGTGCAACCAGGCG |

**Supplementary Table S3.**
Primers used for verification of *clb* genes.

| Gene | Primer | Sequence |
|---|---|---|
| *clbA* | Forward *clbA* | TTTAGGGGTGATGAGTGGAGAGGCT |
| | Reverse *clbA* | TCATCAAACCAGTAGAGATAACTTCCTTCACT |
| *clbB* | Forward *clbB* | TGTTCCGTTTTGTGTGGTTTCAGCG |
| | Reverse *clbB* | GTGCGCTGACCATTGAAGATTTCCG |
| *clbC* | Forward *clbC* | TTGACGGAGGCGTTCGATACTTCAC |
| | Reverse *clbC* | ACTTGTATCACTCGGCGGCAATCAA |
| *clbD* | Forward *clbD* | CGGAGAATGTAGTCGGCGTCCATTT |
| | Reverse *clbD* | CCCTGATTTCACGCCCAAATACCCT |
| *clbF* | Forward *clbF* | CGATTGCCCTCACAGAGCCGAATAT |
| | Reverse *clbF* | AATGCCATGAGAAAATAACCGCCGC |
| *clbG* | Forward *clbG* | CGAATATGCTGCGCTGACCTGTAGT |
| | Reverse *clbG* | ATAGCGATTCTCCAGCAGCAGGTT |
| *clbH* | Forward *clbH* | CTTTGTCGAGTTGCCGGAATACCCT |
| | Reverse *clbH* | TGTGTCTGATCTCCTGTGGTCCCTT |
| *clbI* | Forward *clbI* | TTGAGAATGTACGACTGAACCCGCC |
| | Reverse *clbI* | AATGAATGTCCGCCAGCTTCGAAGA |
| *clbJ* | Forward *clbJ* | TGGCCTGTATTGAAAGAGCACCGTT |
| | Reverse *clbJ* | AATGGGAACGGTTGATGACGATGCT |
| *clbK* | Forward *clbK* | TTGATGATCACCACGCCAGCTTCTT |
| | Reverse *clbK* | GCGGATGGCGGTAGTGATAAGCTAG |
| *clbL* | Forward *clbL* | CACAGGTGTCTATGCCCATCGTTGT |
| | Reverse *clbL* | GCCGACCACTGAGTTTGACTGCTAT |
| *clbM* | Forward *clbM* | TGTTTCAAGGCGCGGGTAAGATCAT |
| | Reverse *clbM* | TAGTCACTCACGGCAACAACACGAG |
| *clbN* | Forward *clbN* | GGCATTCAGTTCGGGTATGTGTGGA |
| | Reverse *clbN* | AACAGAGCTGCCGTAAAGACTCGAC |
| *clbO* | Forward *clbO* | AAGGAGGTGCGGTAAATAACGACGG |
| | Reverse *clbO* | CGGTGGCATGGATCCTTTTCGTTTG |
| *clbP* | Forward *clbP* | CTTGCCGCAGACAATCGTATCCTCT |
| | Reverse *clbP* | CCTGGAGATAGTATACCCGGTGCGA |
| *clbQ* | Forward *clbQ* | CTGTGTCTTACGATGGTGGATGCCG |
| | Reverse *clbQ* | GCATTACCAGATTGTCAGCATCGCC |
