## Supplementary Figures for "SOCS1 expression in prostate epithelial cells is essential for tissue homeostasis and tumor suppression"

Ihsan et al.,

**Supplementary Figures**

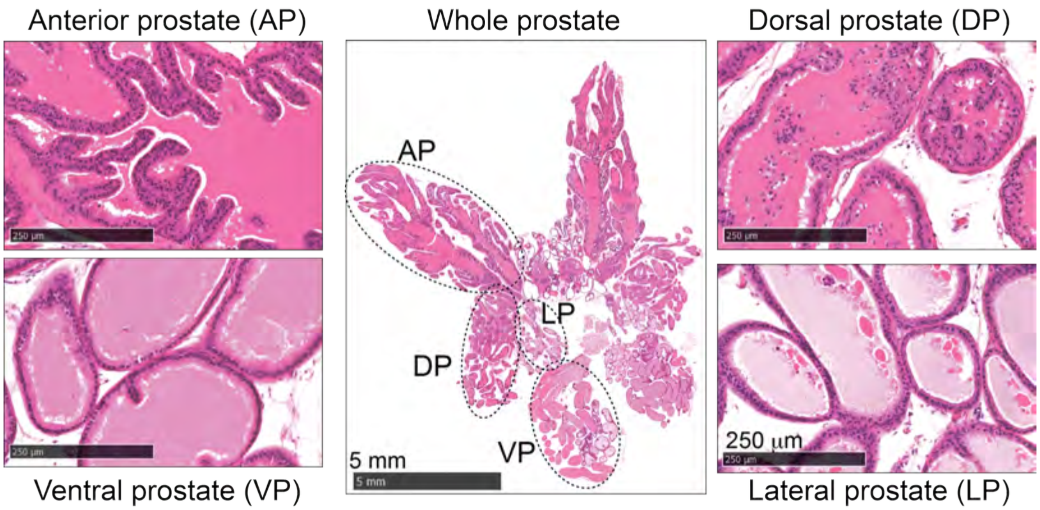


**Supplementary Fig. S1. Whole prostate mount for histological evaluation**. Formalin fixed mouse prostate glands were positioned during paraffin embedding in such a way to reveal all four prostate lobes in one plane of sectioning. Representative hematoxylin and eosin-stained section from one of the best mounts clearly revealing anterior (AP), dorsal (DP), lateral (LP) and ventral (VP) prostate lobes is shown in the centre image. Shown around are representative magnified images of individual lobes, revealing the distinctive features that characterize the glandular acini of each lobe.

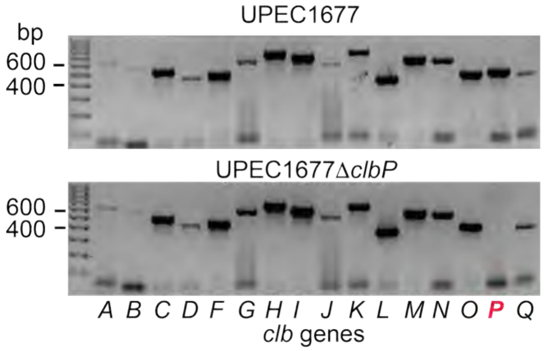


**Supplementary Fig. S2. PCR verification of *clbP* deletion in UPEC1677.** Bacterial DNA was isolated and the intactness of *clb* genes was verified by PCR using primers listed in Supplementary Table S3.

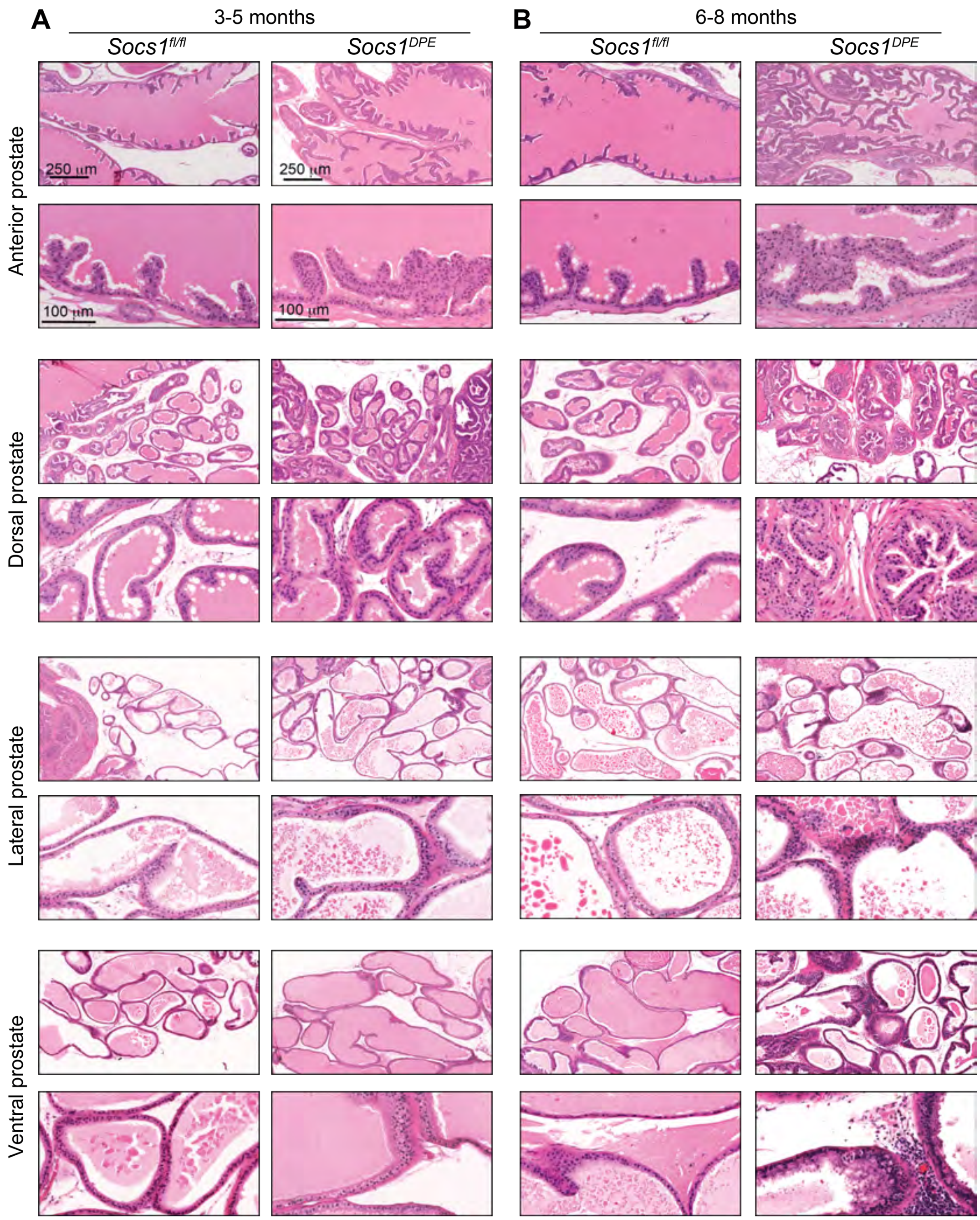


**Supplementary Fig. S3. Progressive development of hyperplastic epithelia in SOCS1-deficient prostate gland**. FFPE-sections of prostate glands from *Socs1*$^{fl/fl}$ and *Socs1*$^{\Delta PE}$ mice of 3-5 months-old (A) and 6-8 months-old (B) age groups were stained with hematoxylin and eosin. Representative regions of anterior, dorsal, lateral and ventral prostate lobes are shown.

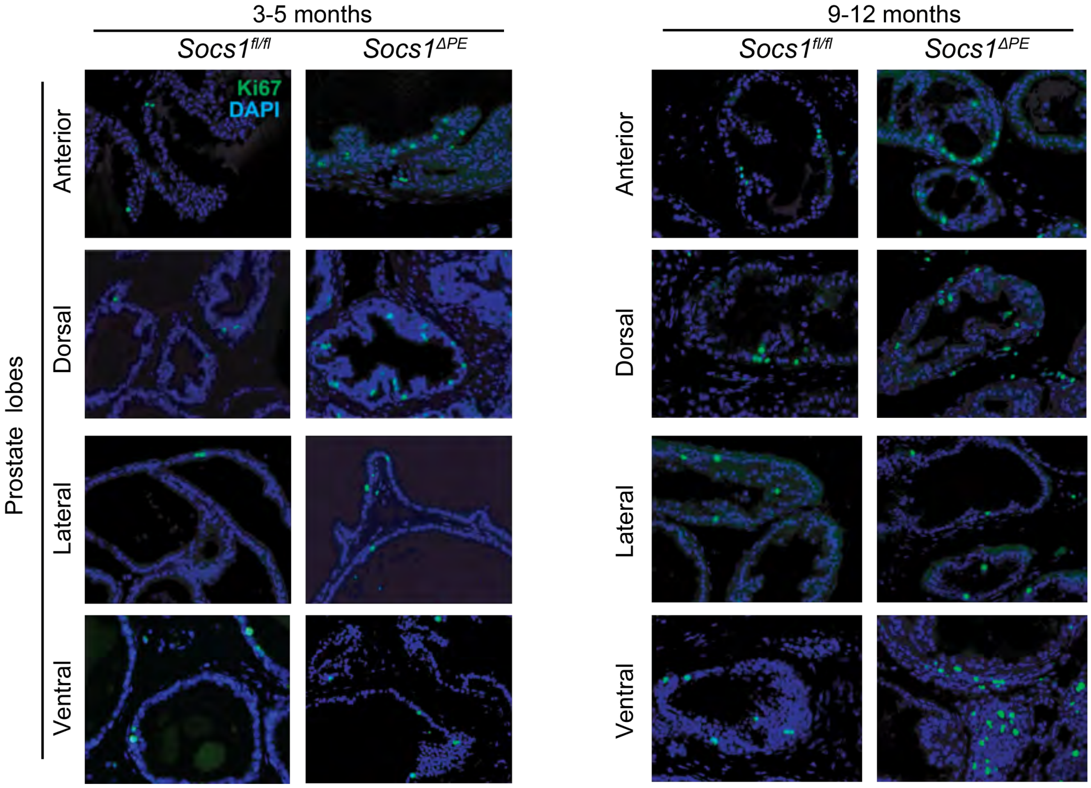


**Supplementary Fig. S4. Increased prostate epithelial cell proliferation in younger and older *Socs1$^{\Delta PE}$* mice**. FFPE-sections of prostate glands from *Socs1$^{fl/fl}$* and *Socs1$^{\Delta PE}$* mice of 3-5 months-old (A) and 9-12 months-old (B) age groups were labelled Ki67 antibody and detected by immunofluorescence. Representative regions of anterior, dorsal, lateral and ventral prostate lobes are shown.

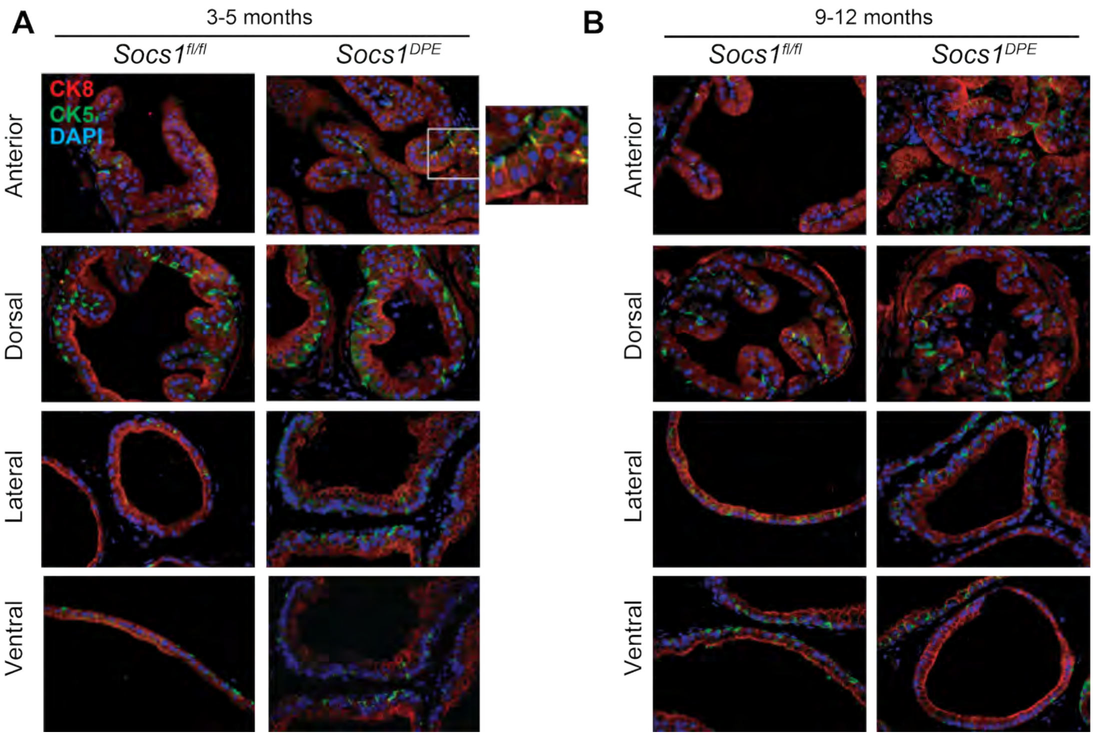


**Supplementary Fig. S5. Distribution of Increased prostate epithelial cell proliferation in younger and older *Socs1$^{\Delta PE}$* mice**. FFPE-sections of prostate glands from *Socs1$^{fl/fl}$* and *Socs1$^{\Delta PE}$* mice of 3-5 months-old (A) and 9-12 months-old (B) were stained for luminal (CK8) and basal (CK5) epithelial cell markers. Representative staining patterns in the prostate lobes of 6-8 months-old mice are shown. An area of the anterior lobe of *Socs1$^{\Delta PE}$* mice containing CK8+CK5+ cells are enlarged to show the double positive cells. (D) Androgen receptor staining in 6-8 months old *Socs1$^{fl/fl}$* and *Socs1$^{\Delta PE}$* mice.

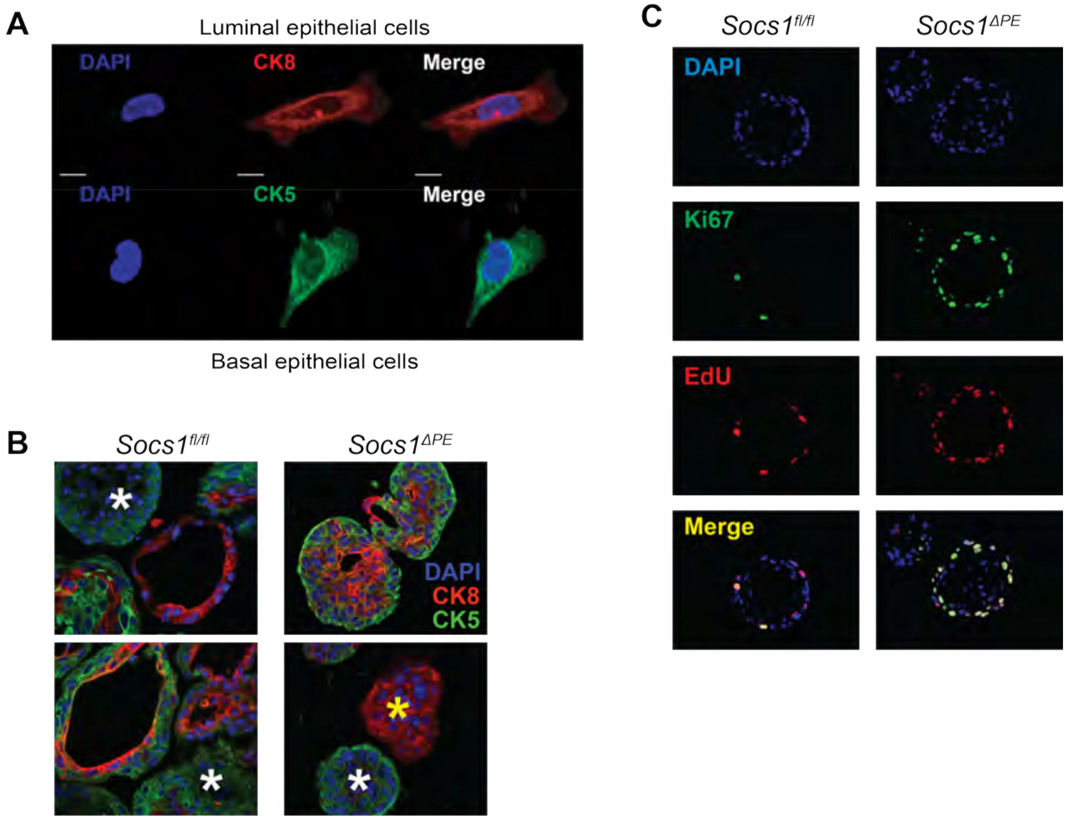


**Supplementary Fig. S6. Generation and characterization of SOCS1 deficient and control prostate organoids.** (A) Single cell suspensions from the prostate gland were stained for luminal (CK8) and basal (CK5) epithelial cell markers, and alpha smooth muscle actin (αSMA) to detect stromal cells. Most cells were positive for CK8 and a few for CK5, and no αSMA+ cells were detected. (B) Additional images of prostate organoids stained for CK8 and CK5 revealing organoids containing only luminal (yellow asterisk) or basal (white asterisks) epithelial cells from both *Socs1$^{fl/fl}$* and *Socs1$^{\Delta PE}$* mice. (C) Representative Ki67 and EdU staining showing increased cell proliferation in SOCS1-deficient prostate organoids.

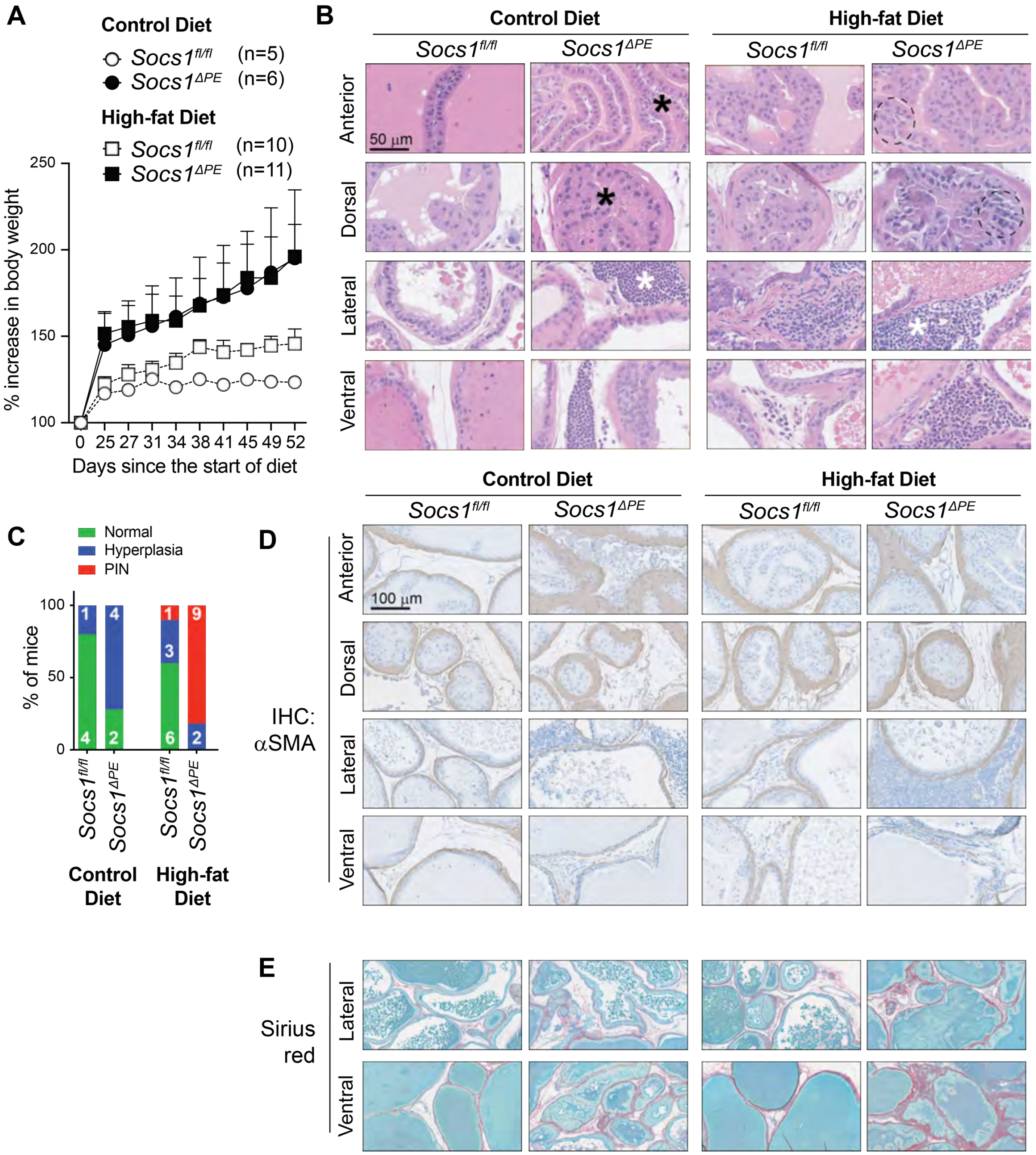


**Supplementary Fig. S7**. Two month-old *Socs1*$^{fl/fl}$ and *Socs1*$^{\Delta PE}$ mice were fed with high-fat diet (HFD) or normal control diet (CD) for 10 months. (A) Gain in body weight during the initial 16 weeks of diet regimen. Number of mice per group is indicated (n). Tukey multiple comparison test was used to compare mice fed with CD or HFD. (B) H&E-stained prostate sections in the different prostate lobes of representative mice, highlighting notable histological features of hyperplasia (black asterisks), prostate intraepithelial neoplasia (PIN)-like lesions (circled) and prominent inflammatory cell infiltration (white asterisks). (C) Cumulative data of hyperplasia and PIN in all groups of mice from anterior and dorsal prostates. Statistics: Chi-Square test. (D) Immunohistochemical staining for alpha smooth muscle actin in representative prostate lobes. (E) Picrosirius red staining for collagen in representative lateral and ventral prostate lobes.

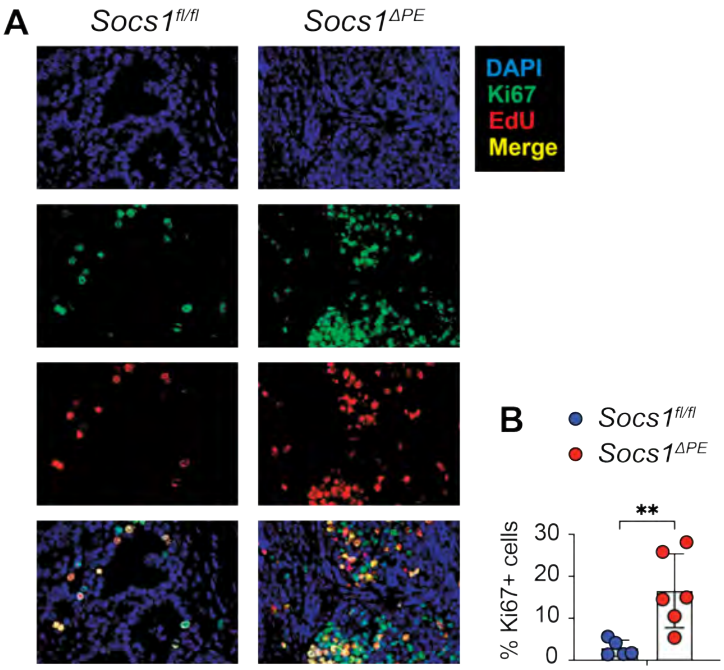


**Supplementary Fig. S8. Increased cell proliferation in the neoplastic prostate tissues of UPEC1677-infected *Socs1$^{\Delta PE}$* mice.** FFPE sections of prostate tissues from UPEC1677-infected and EdU-administered (50 µg per gram bodyweight, i.p.) mice were stained for EdU incorporation into replicating DNA using the Click chemistry-based EdU detection reagent. The slides were subsequently stained for Ki67 examined under Zeiss Axioscope 2 fluorescence microscope at 40× magnification. (A) Representative images of prostate sections from *Socs1$^{fl/fl}$* and *Socs1$^{\Delta PE}$* mice. (B) Quantification of Ki67+ cells from total of ten random fields from 5-6 mice per group. Mean + standard error of mean; Unpaired *t* test. *p* value: ** <0.01.

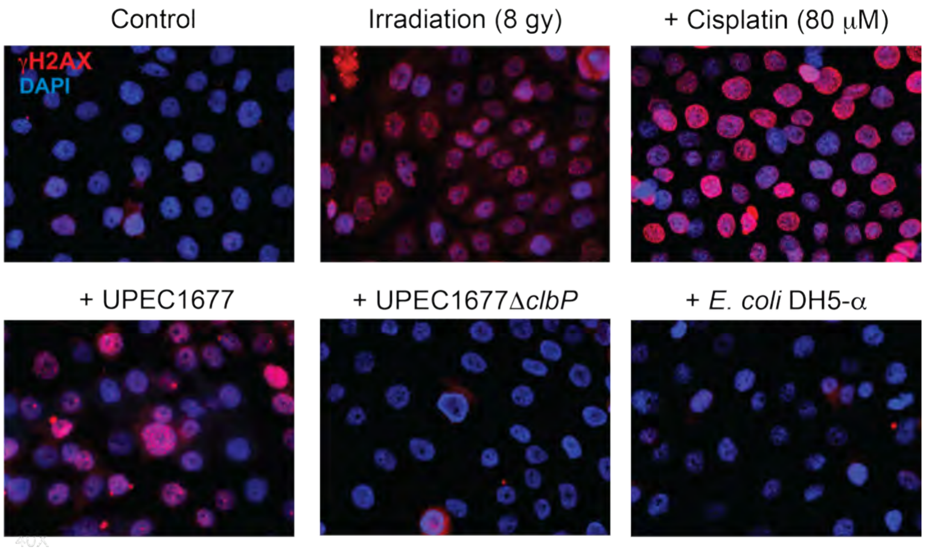


**Supplementary Fig. S9. Deletion of the *clbP* gene in UPEC abolishes its ability to induce DNA damage.** The human benign prostate hyperplasia cell line BHP-1 (purchased from Millipore Sigma Cat# SCC256) and grown on coverslips in RPMI-1640 medium containing 10% FBS were irradiated, or exposed to cisplatin, UPEC1677, UPEC1677*ΔclbP* or *E. coli* DH5a (all bacteria at the multiplicity of infection 200). Infected cultures were washed off bacteria 30 min later in gentamycin (50 µg per mL) containing medium and incubated for further18 h in antibiotic-containing (5 µg/mL gentamicin) medium. The coverslips were fixed, stained with γH2AX antibody and DAPI and examined under a fluorescence microscope.

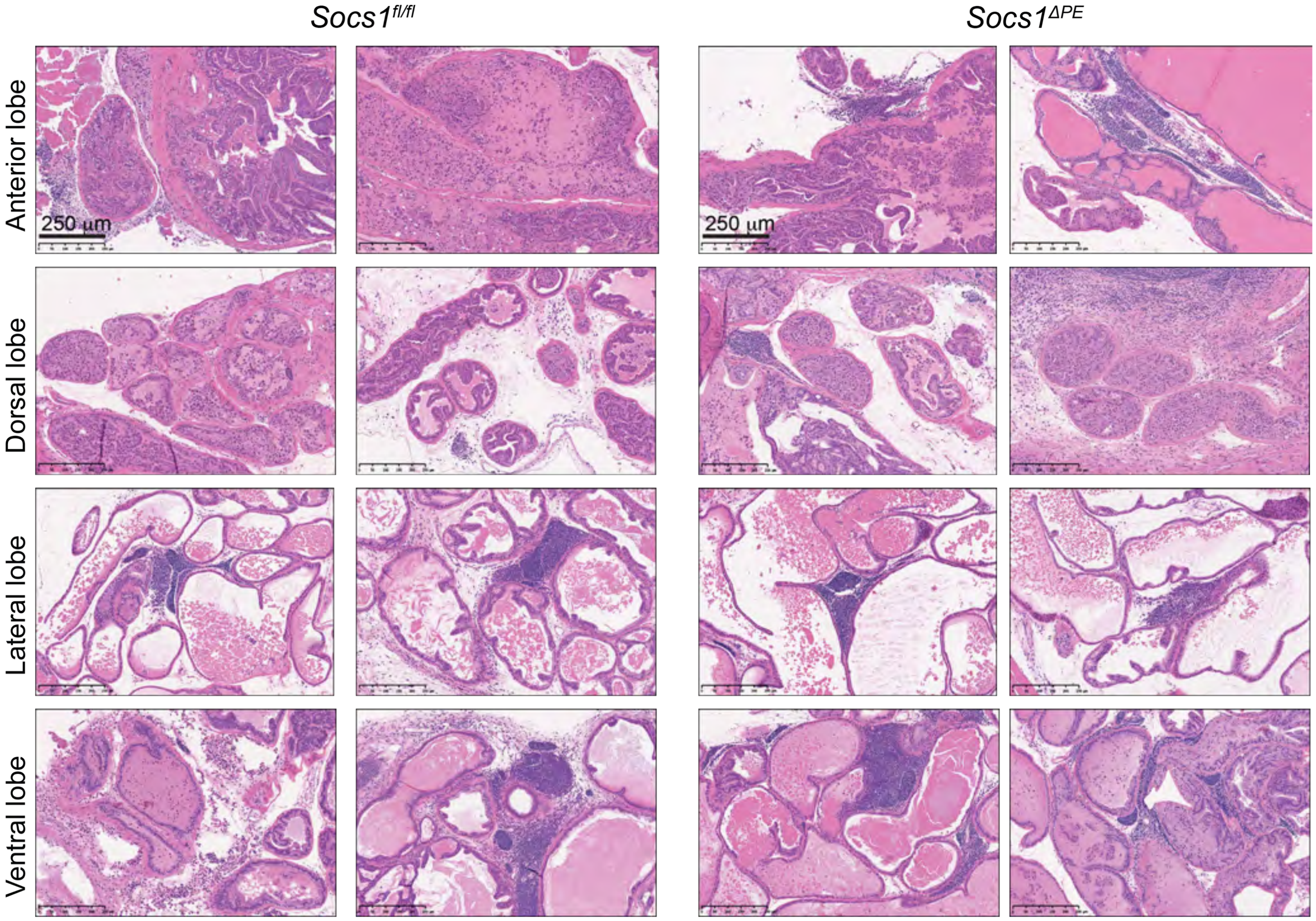


**Supplementary Fig. S10. Immune cell infiltration and hyperplastic changes in the prostates of UPEC1677*ΔclbP*-infected mice.** Prostates of 4 month-old *Socs1*$^{fl/fl}$ and *Socs1*$^{ΔPE}$ mice were inoculated with UPEC1677*ΔclbP* and their prostate glands were examined 8 months later. Hematoxylin and eosin-stained sections showing anterior, dorsal, lateral and ventral prostate lobes of UPEC1677*ΔclbP*-infected *Socs1*$^{fl/fl}$ and *Socs1*$^{ΔPE}$ mice. Representative images of two additional mice for each genotype (other than the ones shown in Fig. 6E) are shown here (scale bar 250 μm).

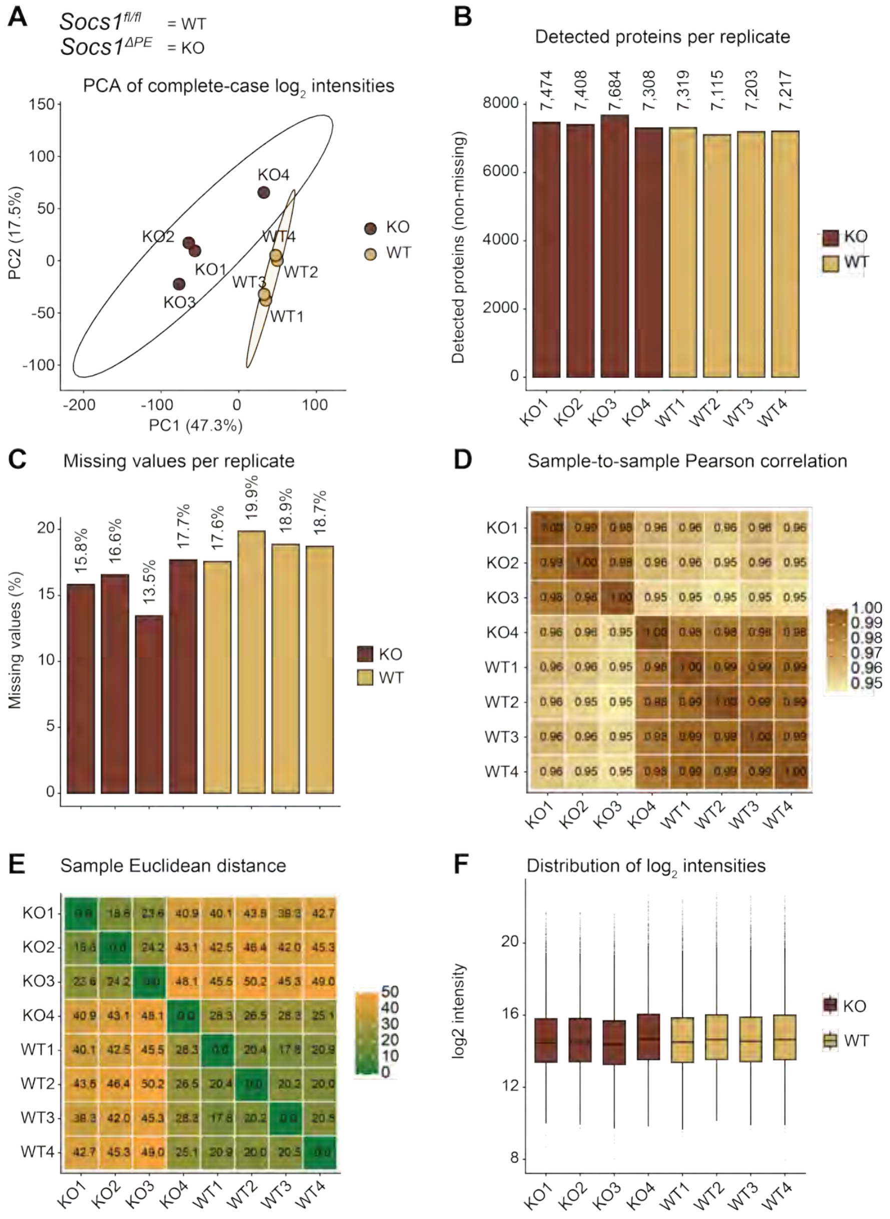


**Supplementary Fig. S11. Evaluation of label-free quantification data for quality control**. (A) Principal component analysis (PCA)of log-transformed intensities for complete-case datasets, where there is a distinct clustering of *Socs1*$^{fl/fl}$ (wild-type, WT) versus *Socs1*$^{\Delta PE}$ (knockout, KO) samples according to the first principal component axis, suggesting condition-specific changes in the proteome with good within-condition clustering. (B) Number of proteins identified in each replicate sample, showing similar numbers of identified proteins in both conditions without large differences. (C) Proportion of missing data in each replicate, indicating similar levels of missing data between the WT and KO conditions. (D) Pearson correlation coefficient matrix of log-transformed intensities, which suggests very high correlation coefficients ($r \approx 0.95$-$1.00$) between replicates for each condition. (E) Heatmap of Euclidean distances between samples, which confirms clustering according to biological condition and showing smaller intra-condition distances compared to inter-condition distances. (F) Histograms of log-transformed intensities between replicates, suggesting consistent distribution of intensities and lack of systematic bias in the data.

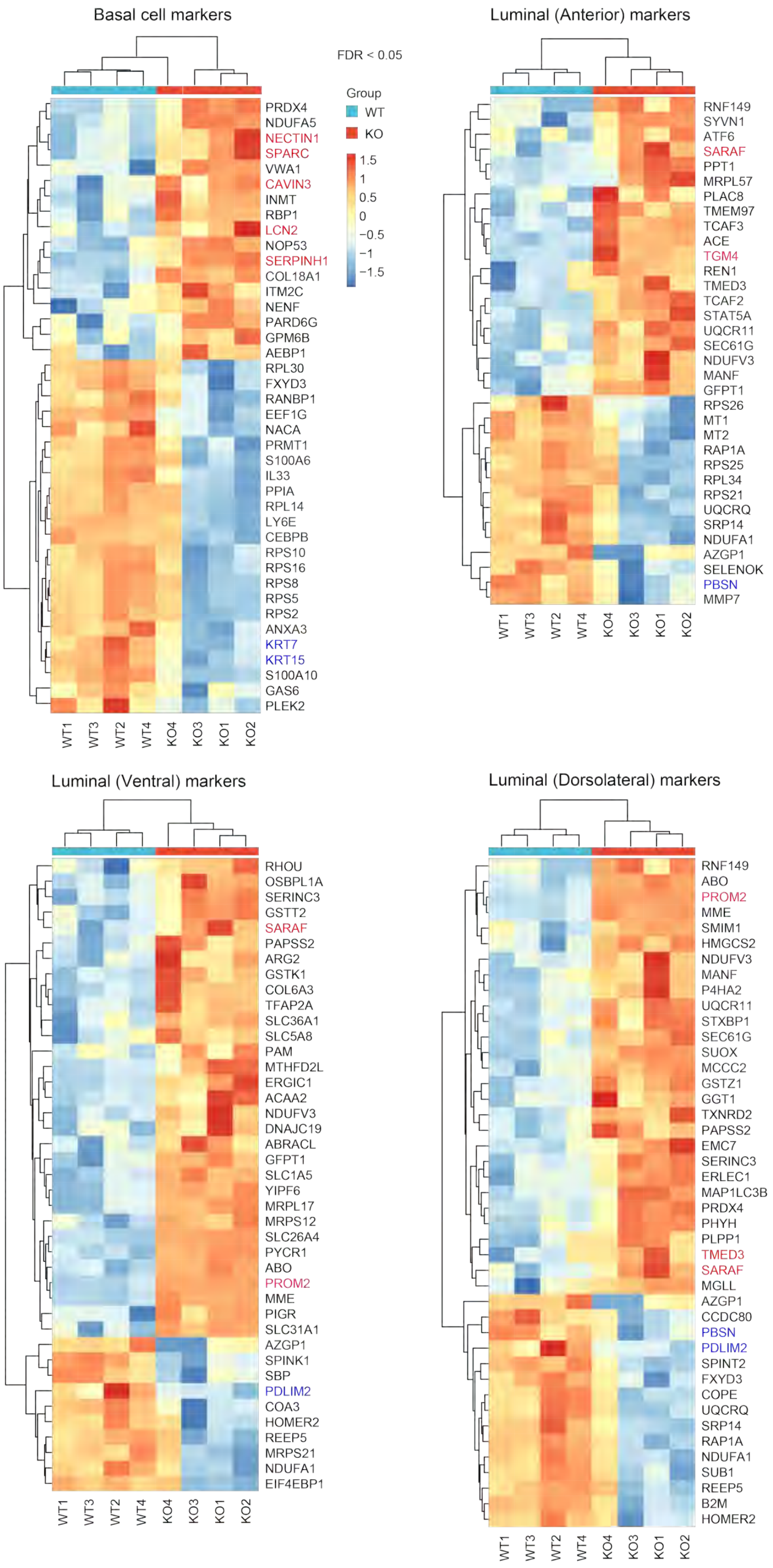


**Supplementary Fig. S12. Modulation of epithelial cell marker expression in SOCS1-deficient prostate epithelial organoids.** Expression levels of proteins associated with differential gene expression signatures of basal epithelial cells (A) and luminal epithelial cells of anterior (B), dorsolateral (C) and ventral (D) lobes of the mouse prostate gland (Ref: 67, 68). Unsupervised clustering analysis of the expression levels of top upregulated and downregulated proteins in SOCS1-deficient organoids compared to control organoids. FDR values are significantly different for all four cell lineage markers ($q$ <0.05).

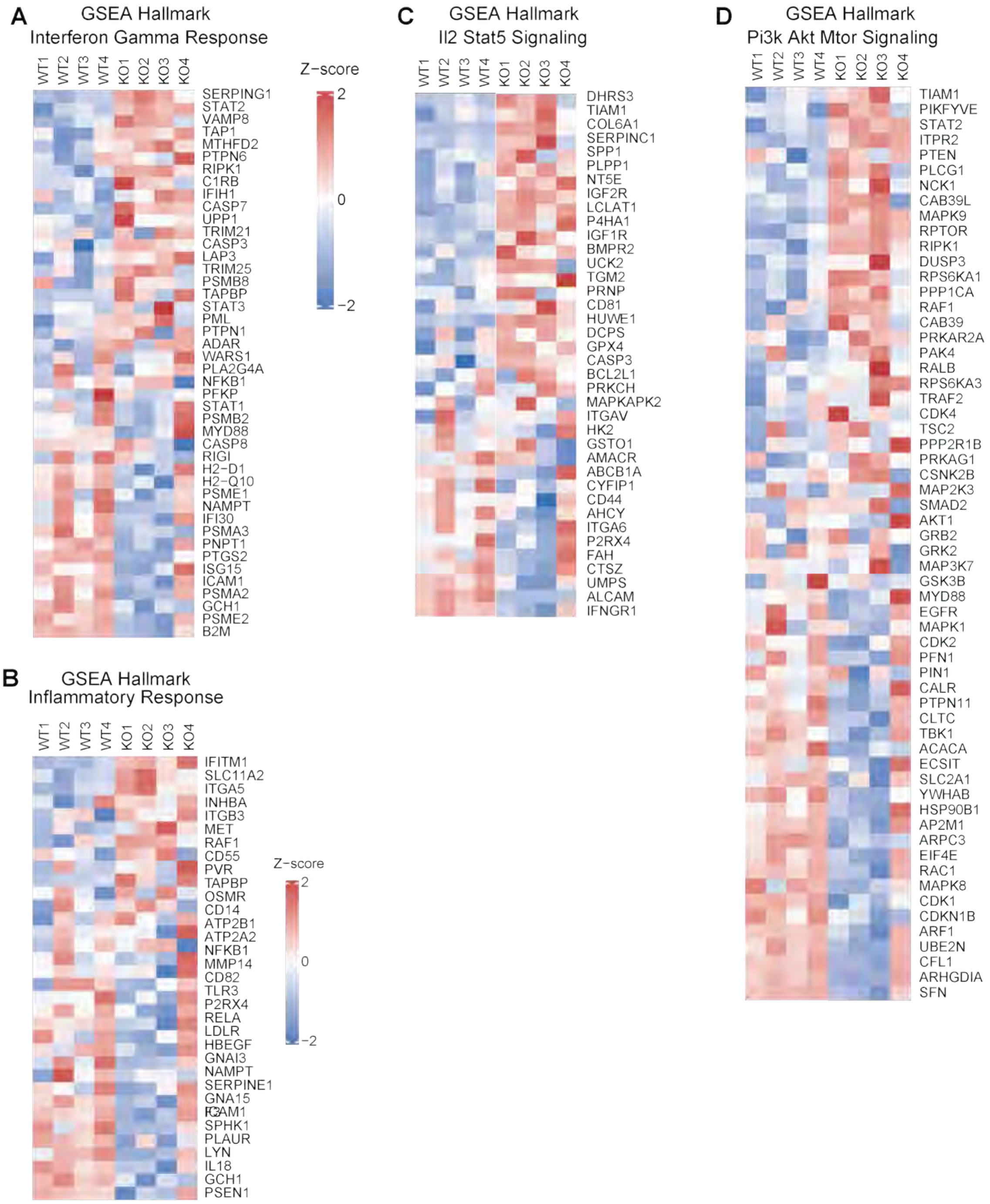


**Supplementary Fig. S13. Heatmap analysis of differentially expressed proteins in SOCS1-deficient prostate epithelial cells within positively enriched GSEA Hallmark pathways.**

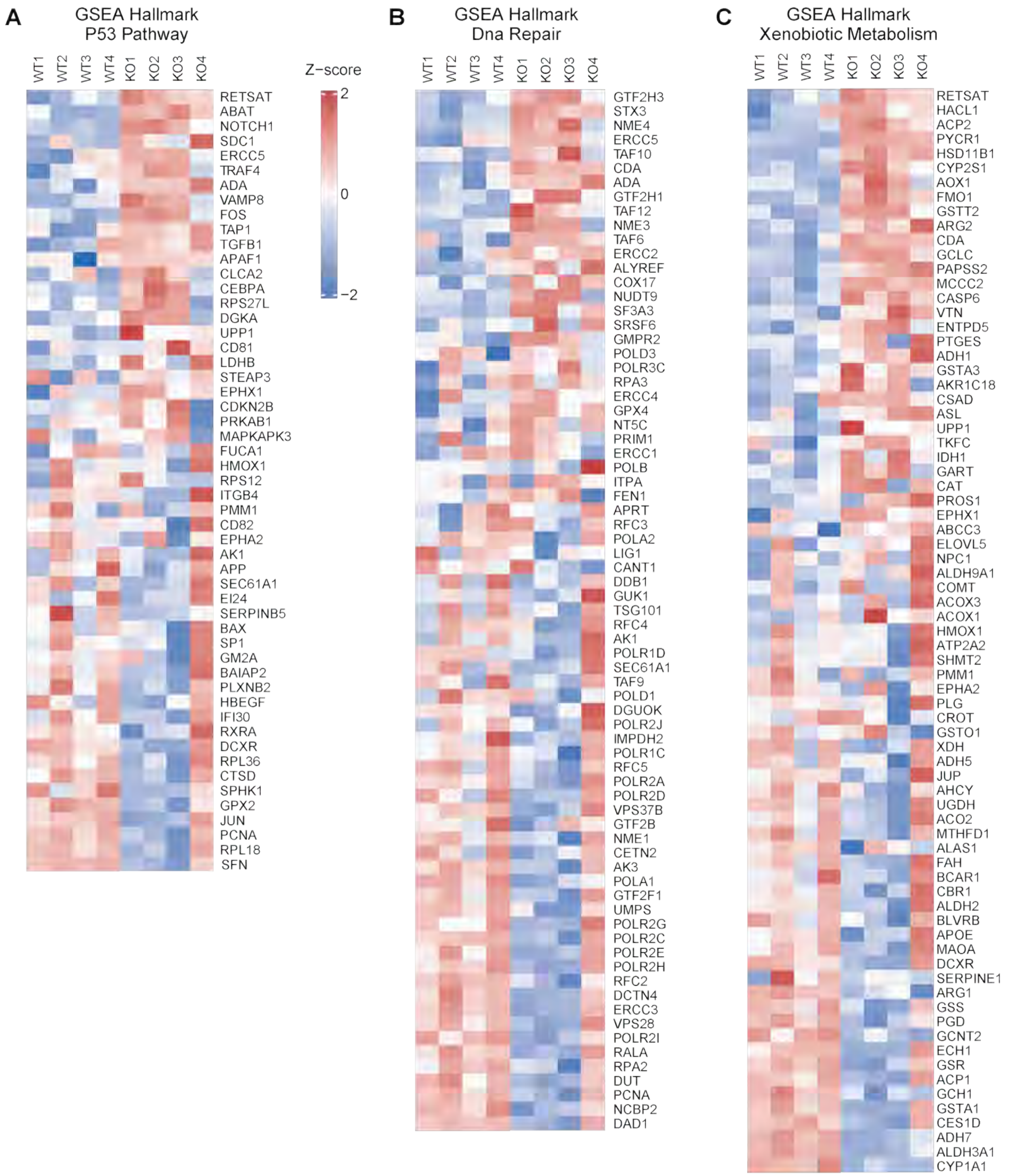


**Supplementary Fig. S14. Heatmap analysis of differentially expressed proteins in SOCS1-deficient prostate epithelial cells within negatively enriched GSEA Hallmark pathways.**
