## Supplementary Methods for "SOCS1 expression in prostate epithelial cells is essential for tissue homeostasis and tumor suppression"

Ihsan et al.,

### Supplementary Methods

#### High-fat diet induced obesity

Eight to twelve week-old *Socs1*$^{fl/fl}$ and *Socs1*$^{\Delta PE}$ mice were fed *ad libitum* a high-fat diet (HFD) with 60% energy from fat (*Research Diets Inc.,* New Brunswick, NJ, USA, Cat # D12492) and matched control diet (CD; Cat #D12450J) with only 10% energy from fat. Mice from all four groups were sacrificed after 9 months of diet regimen and prostate tissues were fixed in 4% PFA and paraffin embedded for histology analysis.

#### EdU labelling *in vitro* and *in vivo*

To label proliferating cells in organoid cultures, 10µM of the nucleoside analog of thymidine, EdU (5-ethynyl-2'-deoxyuridine; ThermoFisher, Cat# C10337) to organoid cultures 8 h before harvesting and processed for fixation and paraffin embedding. To label proliferating cells in prostate tissues of UPEC1677-infected mice (n=5), EdU (50 µg per gram bodyweight; i.p.) 8 h before euthanasia. Prostate tissues were fixed and paraffin embedded. Prostate organoid and tissue sections were deparaffinized followed by antigen retrieval and incubation in blocking buffer. The sections were covered with Click-iT™ EdU detection reagent containing Alexaflour-488 for 1 h at room temperature in the dark and then washed with PBS. Following incubation with primary and secondary antibodies for Ki67 detection, slides were mounted for observation under Zeiss Axioscope 2 fluorescence microscope.

#### Proteomic analysis

#### Protein preparation and protease digestion

Prostate epithelial organoids established from *Socs1*$^{fl/fl}$ (WT) and *Socs1*$^{\Delta PE}$ (KO) mice were collected on day 8, washed with ice-cold PBS to release the organoids from Matrigel and centrifuged. Four biological replicated collected from different experiments were used. The pellets were resuspended in

TrypLE and incubated at 37 °C for 5–10 min with gentle pipetting until a single-cell suspension was obtained. The cells were snap frozen in liquid nitrogen and stored at -80°C. For protein extraction, EasyPep™ Mini MS Sample Prep Kit (Thermo Fisher Scientific, Cat# A40006) was used according to its instructions. Briefly, the resulting epithelial cell pellets were lysed in EasyPep lysis buffer supplemented with universal nuclease (1 µL per 100 µL lysis buffer) to reduce nucleic acid-associated viscosity. Samples were homogenized by Branson Sonifier 150 (Branson Ultrasonics™, USA) and pipetting until a uniform lysate was obtained. Protein concentration was determined using the Pierce BCA Protein Assay Kit (Thermo Fisher Scientific, Cat# 23225) according to the protocol. Briefly, a working reagent was prepared by mixing reagent A and reagent B at a 50:1 ratio, and pre-diluted protein assay standards: Bovine serum albomine (BSA) set (Thermo Fisher Scientific, Cat# 23208) were used to generate a standard curve. Aliquots of samples and standards were incubated with the working reagent at 37 °C for 30 minutes, and absorbance was measured at 562 nm using a microplate reader (Spectrostar Nano, BMG Labtech, Germany). Protein concentrations were calculated based on the standard curve, and samples were normalized accordingly. For each sample, 10 to 100 µg of total protein was transferred to a low-binding microcentrifuge tube and adjusted to a final volume of 100 µL with lysis buffer. Proteins were reduced and alkylated by sequential addition of 50 µL reduction solution and 50 µL alkylation solution, followed by incubation at 95 °C for 10 minutes to ensure complete denaturation and cysteine modification. Samples were cooled to room temperature and digested using a trypsin/Lys-C protease mix (50 µL per sample) at 37 °C for 1-3 hours with shaking to generate MS-compatible peptides. Digestion was terminated by addition of 50 µL digestion stop solution. Peptides were purified using peptide clean-up columns supplied with the kit. Briefly, columns were equilibrated and samples were loaded, followed by sequential washing steps to remove salts, detergents, and other contaminants. Peptides were eluted using elution buffer, collected in low-binding tubes, and dried using a Vacufuge Plus centrifuge concentrator (Eppendorf, Cat# 022820168). Dried peptides were resuspended in 0.1% formic acid in water. Peptide quantification was performed using a NanoDrop 2000/2000c spectrophotometer (Thermo Fisher Scientific) and, if necessary, the concentration was adjusted prior to loading liquid chromatography–mass spectrometry (LC-MS) analysis.

**Liquid chromatography-tandem mass spectrometry (LC-MS/MS)**

Concentrated peptides (250 ng) were separated on a nanoHPLC system (nanoElute, Bruker Daltonics). The samples were loaded onto an Acclaim PepMap100 C18 Trap Column (0.3 mm id x 5 mm, Dionex Corporation) at 4 µL/min consistent flow and peptides were eluted onto a PepMap C18 analytical nanocolumn (1.9 µm beads size, 75 µm x 25 cm, PepSep) heated at 50°C. Peptides were eluted with solvent B (100% ACN & 0.1% FA) in a 5-37% linear gradient with a flowrate of 400 nL/min for ~2 h. The HPLC system was coupled to an TimsTOF Pro ion mobility mass spectrometer containing Captive Spray nano electrospray source (Bruker Daltonics). Data acquisition was done using diaPASEF mode. For each individual Trapped Ion Mobility Spectrometry (TIMS) measurement in diaPASEF mode, a single mobility window consisting of 27 mass steps (with m/z ranging from 114 to 1414 and a mass width of 50 Da) was employed per cycle, which had a 1.27-second duty cycle. This process involves scanning the diagonal line in the m/z-ion mobility plane for +2 and +3 charged peptides.

**Protein identification**

Peptide mass spectra were analyzed using the DIA-NN , an open-source software [1] suite for DIA/ SWATH data processing (https://github.com/vdemichev/DiaNN, version2.5.1 ), installed in an Apptainer container (https://apptainer.org/, version 1.3.5) using docker image provided on the docker hub [2]. Analysis was performed using default parameters except for these options: 2 missed -cleavages was allowed; trypsin digestion was performed for K/R; protein N-term methionine excision as variable modification for the *in-silico* digest. The *Mus musculus* reference proteome UP000000589 was downloaded from the Uniprot website (https://www.uniprot.org/proteomes/UP000000589). The reference proteome contained a total of 63367 proteins. For the FASTA search, DIA-NN was instructed to perform an *in-silico* digest of the sequence database. A mass tolerance accuracy of MS1 and MS2 of 20 ppm was used for precursor and fragment ions, respectively. Minimum and maximum were set for peptide length (7-30 amino acids), and precursor charge (1-5), precursor m/z (100-1700) and fragmentation m/z (100-1500) for *in silico* library generation or library-free search. For the reanalysis, MBR (match between run) was enabled and chosen the smart profiling when creating a spectral library from DIA data. Carboxyamidomethylation (unimod4), and oxidation (M) (unimod35) were set as fixed modifications and N-terminal protein acetylation was set as a variable modification. DIANN protein group matrix was filtered using custom Perl script to extract protein groups with a single protein.

**Data analysis**

R (v4.5.0) was used to analyze protein intensity data. Data were log2-transformed (log2[x + 1]) and filtered to retain proteins with valid gene symbols. The limma package [3,4] was used to evaluate differential expression between conditions (KO versus WT [$Socs1^{\Delta PE}$ versus $Socs1^{fl/fl}$]), and ranked gene lists were produced using moderated *t*-statistics. Gene set enrichment analysis (GSEA) was performed using gseKEGG for unbiased pathway analysis [5,6]. KEGG pathways were retrieved using KEGGREST and mapped using the org.Mm.eg.db package (Release 3.22). To maintain biological ordering, ComplexHeatmap was used to create heatmaps based on row-wise Z-score normalization of either replicate-level or group-averaged data (KO, WT) without clustering. Using ggplot2 package, GSEA plots were created that showed ranked statistics, gene distribution, enrichment score profiles and highlighted genes [7]. For lineage markers, differential expression and FDR calculations were performed separately within the published marker list [8,9] instead of against the full proteome. Next, proteins with fewer than two valid values per group were excluded to avoid making unreliable estimates, and outputs were plotted based on both adjusted p-values (FDR) and raw p-values. For visualization, expression values were log2-transformed and row-scaled (z-score normalization).

**Data availability**

Curated mass spectrometry data of intensity profiles of proteins detected in all biological replicates, and expression levels of epithelial lineage markers are given in supplementary spreadsheets.
